# TRF1 and TRF2 form distinct shelterin subcomplexes at telomeres

**DOI:** 10.1101/2024.12.23.630076

**Authors:** Tomáš Janovič, Gloria I. Perez, Jens C. Schmidt

**Affiliations:** Institute for Quantitative Health Science and Engineering, Michigan State University, East Lansing; Department of Obstetrics, Gynecology and Reproductive Biology, Michigan State University, East Lansing

## Abstract

The shelterin complex protects chromosome ends from the DNA damage repair machinery and regulates telomerase access to telomeres. Shelterin is composed of six proteins (TRF1, TRF2, TIN2, TPP1, POT1 and RAP1) that can assemble into various subcomplexes *in vitro*. However, the stoichiometry of the shelterin complex and its dynamic association with telomeres in cells is poorly defined. To quantitatively analyze the shelterin function in living cells we generated a panel of cancer cell lines expressing HaloTagged shelterin proteins from their endogenous loci. We systematically determined the total cellular abundance and telomeric copy number of each shelterin subunit, demonstrating that the shelterin proteins are present at telomeres in equal numbers. In addition, we used single-molecule live-cell imaging to analyze the dynamics of shelterin protein association with telomeres. Our results demonstrate that TRF1-TIN2-TPP1-POT1 and TRF2-RAP1 form distinct subcomplexes that occupy non-overlapping binding sites on telomeric chromatin. TRF1-TIN2-TPP1-POT1 tightly associates with chromatin, while TRF2-RAP1 binding to telomeres is more dynamic, allowing it to recruit a variety of co-factors to chromatin to protect chromosome ends from DNA repair factors. In total, our work provides critical mechanistic insight into how the shelterin proteins carry out multiple essential functions in telomere maintenance and significantly advances our understanding of macromolecular structure of telomeric chromatin.

## INTRODUCTION

Telomeres are essential for genome stability, serving as protective caps at the ends of eukaryotic chromosomes. Human telomeres are composed of 2-15 kb of double-stranded (ds) repetitive DNA (TTAGGG) followed by a 3’ single-stranded (ss) overhang, 50-300 nucleotides in length (1). Chromosome ends resemble DNA double-strand breaks and must be protected from inappropriate DNA damage response (DDR) recognition and DNA repair pathway activation. This protection is mediated by the human shelterin complex (Fig. 1A), composed of six subunits (TRF1, TRF2, TIN2, TPP1, POT1, and RAP1), which specifically binds to telomeric DNA and counteracts the recognition of telomeres by the DNA repair machinery (2–4). In addition, shelterin recruits telomerase to telomeres to compensate for incomplete DNA replication of chromosome ends (5). In most cancers (∼90%), telomeres are maintained via telomerase (6,7). However, the remaining 10% of tumors rely on Alternative Lengthening of Telomeres (ALT), a homologous recombination based process unique to cancer cells (8,9). How the shelterin complex carries out these distinct functions, balancing the recruitment of telomerase or proteins involved in ALT, with restriction of access for DNA damage repair proteins remains a critical unaddressed question in the field.

**Figure 1.**
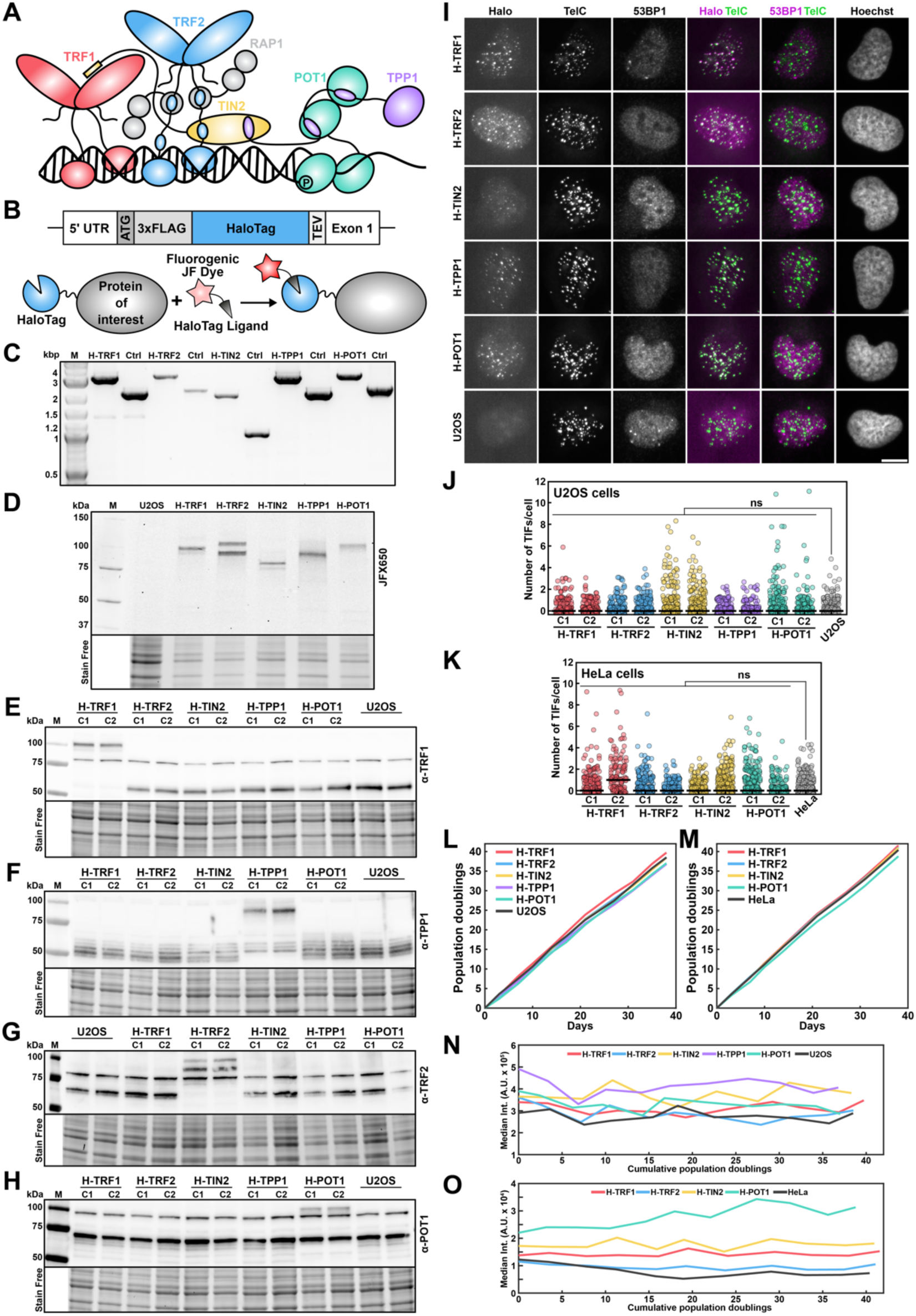
Generation and functional characterization of a panel of cell lines expressing HaloTagged shelterin components. **(A)** Schematic model of the shelterin complex bound to telomeric DNA. **(B)** Design of the N-terminal knock-in of 3xFLAG-HaloTag followed by TEV cleavage site using CRISPR/Cas9 mediated genome editing (top) and the illustration of covalent labeling of HaloTagged protein of interest with Janelia fluor (JF) fluorescent ligands (bottom). **(C)** Genomic PCR result using primers outside of homology arms showing homozygous insertion of HaloTag to shelterin genes in U2OS cells. **(D)** Representative in-gel fluorescence image demonstrating the expression of HaloTagged shelterin proteins at correct size in U2OS cells. **(E-H)** Western blot analysis of U2OS cell lines expressing HaloTagged shelterin proteins using the indicated antibodies confirming protein expression and homozygous insertion. C1 and C2 denote clone number. **(I)** IF-FISH analysis of U2OS cell lines expressing HaloTagged shelterin proteins. The HaloTag was labeled using JFX650 HaloTag-ligand, immunofluorescence against 53BP1 was used to analyze DNA damage signaling at telomeres (represented as TIFs), and PNA FISH to mark telomeres. Scale bar = 10 µm. **(J)** TIFs analysis of U2OS cells expressing HaloTagged shelterin proteins. **(K)** TIFs analysis of HeLa cells expressing HaloTagged shelterin proteins. **(L)** Growth curve of U2OS cells expressing HaloTagged shelterin proteins. **(M)** Growth curve of HeLa cells expressing HaloTagged shelterin proteins. **(N)** Relative telomere length determined using Flow-FISH of U2OS cells expressing HaloTagged shelterin proteins over time. **(O)** Relative telomere length determined using Flow-FISH of HeLa cells expressing HaloTagged shelterin proteins over time.

The shelterin subunits TRF1 and TRF2 are structurally closely related, both forming homodimers mediated by their TRFH-domains, and specifically binding to double-stranded telomeric DNA via C-terminal MYB-domains (10). A major difference between TRF1 and TRF2 lies in a short region of disordered residues at their N-terminus, which in the case of TRF1 is rich in acidic amino acids, while TRF2 contains basic residues (11,12). In addition to mediating dimerization, the TRFH domains of TRF1 and TRF2 facilitate interactions with other critical telomeric proteins. The TRFH domain of TRF1 associates directly with a short peptide in the C-terminus of TIN2, while the same region of TRF2 has been shown to interact with the nuclease Apollo and the endonuclease adaptor SLX4 in addition to binding to the C-terminus of TIN2 (13–17). TRF2 can also associate with the TRFH domain of TIN2 via a short peptide in the hinge region connecting its TRFH and MYB domains (Fig. 1A) (18,19). While TIN2 can associate with TRF2 in two different locations, its recruitment to telomeres in cells is largely mediated by TRF1 (20,21). It is therefore unclear how the interaction between TRF2 and TIN2 observed in *in vitro* reconstitutions and by co-immunoprecipitation contributes to telomere maintenance in cells (21–23).

TRF1 and TRF2 carry out distinct functions at telomeres. TRF2 promotes the formation of telomeric loops (T-loops), where the single-stranded overhang invades the double-stranded telomeric DNA to create a lasso-like structure, which is crucial for preventing the activation of ATM signaling and suppression of non-homologous end joining (NHEJ) (24–28). In contrast, TRF1 facilitates telomere replication and suppresses ATR signaling via the recruitment of TIN2, TPP1, and POT1 (21,29,30). POT1 specifically binds to the single-stranded 3’ overhang with its two OB-fold domains, preventing RPA loading (29,31,32). In addition, POT1 tightly caps the ds-ss telomeric junction by recognizing the 5’-phosophate of the terminal nucleotide using its POT-hole structural feature within the first OB-fold domain (33). Together, these properties of POT1 prevent the activation of ATR signaling at telomeres (4,29). POT1 forms a heterodimer with TPP1, which in turn associates with the TRFH domain of TIN2 via a short peptide in its C-terminus (18,23,34). In addition to tethering POT1 to TIN2, TPP1 is also critical for telomerase recruitment to telomeres (5,35). Thus, TIN2 represents the central hub of the shelterin complex connecting proteins associated with double-stranded telomeric DNA (TRF1, TRF2, RAP1) to proteins bound to the single-stranded overhang (POT1, TPP1) (15,18,36,37). Finally, RAP1 binds to TRF2 and modulates the TRF2’s interaction with telomeric DNA (38,39). The shelterin complex is highly dynamic with extensive conformational variability, which has hindered high-resolution structural analysis of the entire complex, although structures of many individual folded domains of shelterin subunits have been determined (40–42).

*In vitro* reconstitutions have revealed a wide variety of possible shelterin assemblies, ranging from subcomplexes (e.g. TIN2-TPP1-POT1, TRF1 or TRF2-TIN2-TPP1-POT1) to a fully assembled dimeric complex, containing two copies of all six shelterin proteins (41–44). Previous work in human cancer cells has suggested that TPP1 and POT1 are less abundant than TRF1, TRF2, and TIN2 and that the number of shelterin proteins associated with telomeres ranges from 65 molecules per telomere for POT1 to 740 molecules per telomere for TIN2. This uneven stoichiometry of shelterin subunits implies that multiple subcomplexes exist rather than a single six-protein molecular entity (45). However, the exact copy number of shelterin proteins at telomeres has not been directly determined and how it is dynamically regulated to coordinate the different functions of the shelterin complex in telomere maintenance is unknown.

To precisely define the stoichiometry and dynamic association of the shelterin proteins with telomeres, we introduced the HaloTag into the endogenous loci of all six shelterin proteins in telomerase-positive (HeLa) and ALT (U2OS) cells using CRISPR/Cas9 genome editing (46). Using these cell lines, we directly measured the shelterin abundance at telomeres in intact cells. We found that 30-40 copies of each shelterin protein are represented at telomeres in HeLa cells, while the stoichiometry of shelterin at telomeres is altered in U2OS cells. Furthermore, live-cell single-molecule imaging revealed that TRF2 and RAP1 subunits are dynamically associated with chromosome ends, while the rest of the shelterin components are tightly bound to telomeres. In addition, our results indicate that TRF1 and TRF2 occupy distinct binding sites at telomeres. These observations suggest that shelterin exists primarily in two subcomplexes that fulfil distinct functions. The tightly bound TRF1-TIN2-TPP1-POT1 subcomplex that prevents ATR activation, controls telomerase access to chromosome ends and promotes DNA replication, while the dynamic binding of TRF2-RAP1 likely facilitates the recruitment of a variety of factors involved in telomere protection from the DNA repair machinery, including T-loop formation. In total, we define the biochemical and biophysical properties of shelterin proteins in living cells and we provide critical insights into overall telomeric architecture and the molecular mechanism by which shelterin orchestrates multiple essential functions to achieve telomere homeostasis.

## RESULTS

### Generation and functional characterization of a panel of cell lines expressing HaloTagged shelterin components

To analyze stoichiometry and dynamics of the shelterin proteins in living human cells, we generated a panel of telomerase-positive (HeLa-EM2-11ht (47), referred to as HeLa) and ALT (U2OS) cancer cell lines expressing HaloTagged shelterin components from their endogenous loci using CRISPR/Cas9 genome editing (46). We introduced a 3xFLAG-HaloTag (referred to as Halo throughout the manuscript) at the N-terminus of *TRF1*, *TRF2*, *TIN2*, *TPP1*, and *POT1* loci (Fig 1A,B). For the *TRF2* and *TPP1* genes, we targeted the start codons of their major isoforms at amino acids position 42 and 89, respectively. In addition, we targeted the *TIN2* and *TRF2* loci in HeLa 1.3 cell line, which has longer telomeres (∼20 kb) compared to HeLa cells. We confirmed the homozygous insertions by genomic PCR and Sanger sequencing (Fig. 1C, Fig. S1A), and we validated the expression of edited proteins using in-gel fluorescence (Fig. 1D, Fig. S1B) and western blot (Fig. 1E-H, Fig. S1C-E). Importantly, we were able to generate clonal HaloTagged TPP1 cell lines in HeLa cells but they stopped proliferating shortly after clonal isolation (Fig. S1D,E), suggesting that the N-terminal tag on TPP1 affected telomerase recruitment. We also could not validate TIN2 expression by western blotting due to lack of an antibody that could reliably detect TIN2 at endogenous expression levels. However, in-gel fluorescence (Fig. 1D, Fig. S1B) and genomic PCR (Fig. 1C, Fig. S1A) clearly demonstrated the expression of Halo-TIN2 at the expected size and targeting of all alleles, respectively. All tagged shelterin proteins, besides Halo-TIN2, were expressed at similar levels to their untagged counterparts, and HaloTagging the shelterin subunits did not affect the expression levels of the other shelterin components (Fig. 1E-H, Fig. S1C-E). The introduction of HaloTag at the N-terminus of the shelterin proteins allowed us to detect protein isoforms that resulted from alternative splicing downstream of the site of tag insertion. For TRF2 we detected two distinct bands with similar abundance by fluorescence gels in HeLa, HeLa 1.3, and U2OS cells (Fig. 1D, Fig. S1B). Consistent with previous findings, we also observed two TIN2 isoforms (Fig. 1D, Fig. S1B) (48).

To assure that the HaloTag did not affect protein function, we confirmed that the tagged shelterin proteins localized to telomeres and that their expression did not lead to the formation of telomere dysfunction-induced foci (TIF), marked by 53BP1 (Fig. 1I-K, Fig. S1F). To further validate the functionality of tagged shelterin proteins, we cultured cells over a period of 40 days and analyzed their grow rate and used flow cytometry coupled with telomeric PNA FISH (Flow-FISH) to determine their relative telomere length. The results demonstrate that all cell lines had comparable growth rates (Fig. 1L,M), and that telomere length was stable over time in the U2OS cell lines (Fig. 1N). In HeLa cells, telomere length in the Halo-TRF1, TRF2 and TIN2 lines was constant over time (Fig. 1O). In contrast, telomere length in HeLa cells expressing Halo-POT1 increased slightly over the same time period (Fig. 1O). A similar phenotype was observed in cancer cell lines expressing POT1 mutants that alter the DNA binding affinity of POT1 (49). This suggests that the HaloTag on POT1 most likely alters its DNA binding properties. However, HeLa cells expressing Halo-POT1 did not display defect in cell growth (Fig. 1M) or telomere protection (Fig. 1K) and HaloTagging POT1 in U2OS cells did not lead to telomere elongation (Fig. 1N), indicating that this phenotype is telomerase dependent.

Collectively, these results demonstrate that we have successfully generated a panel of telomerase-positive and ALT cancer cell lines expressing HaloTagged shelterin components from their endogenous loci. While the HaloTagged shelterin components are fully functional in chromosome end protection in both cell lines, tagging POT1 in HeLa cells lead to an increase in telomere length.

### Total cellular abundance of shelterin proteins

To quantify the absolute cellular abundance of the tagged shelterin proteins, we used an in-gel fluorescence approach previously described (50,51). The HaloTagged proteins were quantitatively labeled with JF657 HaloTag-ligand, cell lysates were analyzed by SDS-PAGE, and the fluorescence intensity of the HaloTagged shelterin proteins was compared to recombinant HaloTag standard (Fig 2A,B). The HaloTag standard was also supplemented with cell lysate from a known number of U2OS or HeLa cells. Standard curves for both HaloTag and total protein stain signal intensity were linear over a broad dynamic range, which allowed us to quantitatively measure the total cellular abundance of HaloTagged shelterin proteins (Fig. S2A,B). To minimize the impact of protein composition of the HaloTagged shelterin proteins on the SDS-PAGE analysis, we used TEV protease to cleave the HaloTag of the shelterin proteins in cell lysates to correct for differences in the banding pattern (Fig. S2C-E). Our results demonstrate that TRF2 is the most abundant shelterin protein in U2OS (∼60,000 proteins/cell) and HeLa (∼85,000 proteins/cell) cells (Fig. 2C,D). TRF1 is present at ∼10-12,000 proteins/cell in both U2OS and HeLa cells (Fig. 2C,D), while TIN2 and POT1 were more abundant in HeLa cells (∼35,000 proteins/cell) compared to U2OS cells (less than 10,000 proteins/cell for TIN2 and ∼12,000 proteins/cell for POT1) (Fig. 2C,D). TPP1 was expressed at ∼20,000 proteins/cell in U2OS cells (Fig. 2C). In HeLa 1.3 cells, both TRF2 and TIN2 were expressed at ∼40,000 proteins/cell (Fig. S2F,G). While the abundance of shelterin proteins are comparable to those determined by Takai et al. using quantitative western blotting (45), the protein number of TIN2 in U2OS cells presented here is approximately 10-fold lower than previously measured. Since we were unable to verify that Halo-TIN2 is expressed at similar levels to endogenous TIN2 it is possible that we are underestimating the levels of TIN2. Altogether, we have measured the total cellular abundance of the HaloTagged shelterin factors in U2OS and HeLa cells. TRF2 was substantially more abundant than the other shelterin subunits, which could allow TRF2 to recruit a wide variety of factors to telomeres in the context of telomere end protection.

**Figure 2.**
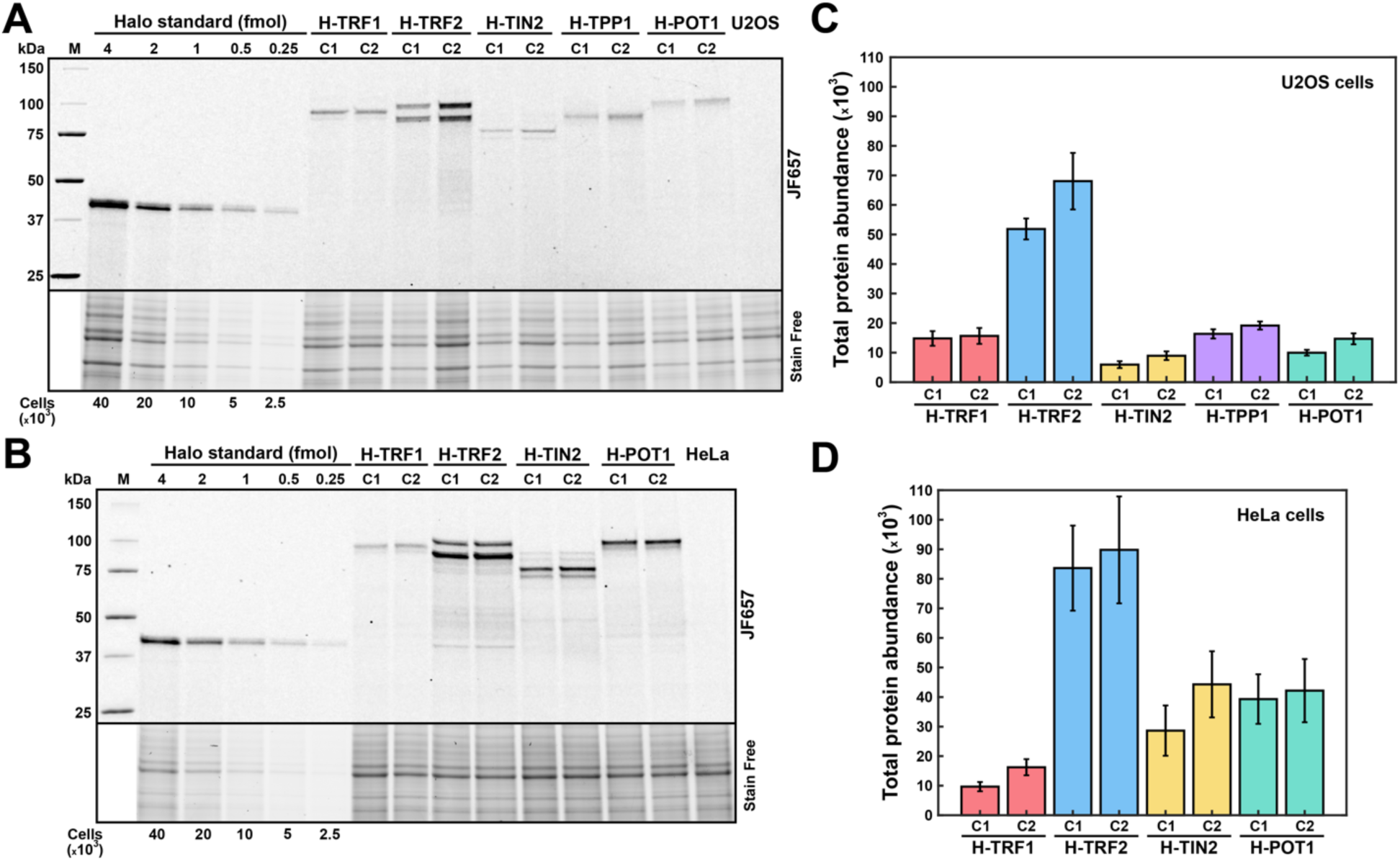
Total cellular abundance of HaloTagged shelterin proteins. **(A-B)** Representative in-gel fluorescence result comparing quantitatively labeled Halo-shelterin proteins in **(A)** U2OS cells and **(B)** HeLa cells with a known amount of recombinant, fluorescently labeled 3x-FLAG-HaloTag standard. **(C-D)** Quantification of total cellular HaloTagged shelterin abundance in **(C)** U2OS cells and **(D)** HeLa cells (*N* = 3, mean ± SD).

### Shelterin abundance and stoichiometry at telomeres

While total cellular abundance of shelterin provides important information regarding overall shelterin stoichiometry, it does not report on the shelterin complex composition at telomeres. To determine the number of shelterin proteins at individual telomeres, we carried out fluorescence photobleaching experiments (Fig. 3A-C, Movie S1). We quantitatively labeled the HaloTagged shelterin proteins with JFX650 HaloTag-ligand, fixed cells, and immediately imaged them using a high laser power to monitor photobleaching of telomeric shelterin signals. Time traces of the fluorescence intensity of telomeric foci allowed us to measure the intensity of single fluorophores by determining the magnitude of individual photobleaching steps, which were clearly discernible in the intensity profiles (Fig. 3B,C). To quantify the telomeric copy number of the HaloTagged shelterin proteins we divided initial Halo-shelterin foci intensity, acquired using identical microscope settings, by the fluorescence intensity of a single-fluorophore. To eliminate the contribution of DNA replication and potential overlap of the telomeres from sister chromatids, cells were synchronized using a double thymidine block at the beginning of S-phase (Fig. S3A). The number of telomeric foci detected per nucleus were slightly lower than the expected number of chromosome ends in the respective cell line (e.g. ∼125 detected foci compared to 140-150 chromosome ends in HeLa cells, Fig. S3B-D). This suggests that a small fraction of telomeric signals are not detected because they are below the detection limit or multiple telomeric signals spatially overlap. The telomeric foci number was lower in U2OS cells (∼100 foci per nucleus, Fig. S3D), suggesting that a larger fraction of telomeres are clustered together, consistent with PML body formation (52). In HeLa cells, the telomeric shelterin abundance ranged from 25-35 copies for TRF1, TRF2, and TIN2 to around 40 copies for POT1 per telomere (Fig. 3D). Importantly, the increased telomere length in the Halo-POT1 HeLa cells (Fig. 1O) compared to the other HeLa cell lines could be responsible for the higher number of POT1 molecules per telomere, compared to TIN2, which is upstream of POT1 in the telomeric recruitment hierarchy (23). Therefore, our results indicate that all shelterin factors are present at telomeres in approximately equal numbers. In HeLa 1.3 cells, TRF2 and TIN2 were present at around 40 copies per telomere (Fig. 3D), even though HeLa 1.3 cells have longer telomeres than HeLa cells (∼20 kbp vs ∼4 kbp, respectively). In U2OS cells, the telomeric copy number of TRF1 and TRF2 (both around 40 copies per telomere) were slightly higher than in HeLa cells, while the number of TIN2 molecules was lower (∼10 copies per telomere). TPP1 was present in a higher copy number (∼60 copies per telomere) than TIN2, while ∼30 copies of POT1 localized to telomeres in U2OS cells. In particular for TRF1, TRF2, and TIN2 the telomeric protein number determined here are substantially lower than previous estimates reported in the literature based on chromatin fractionation (45).

**Figure 3.**
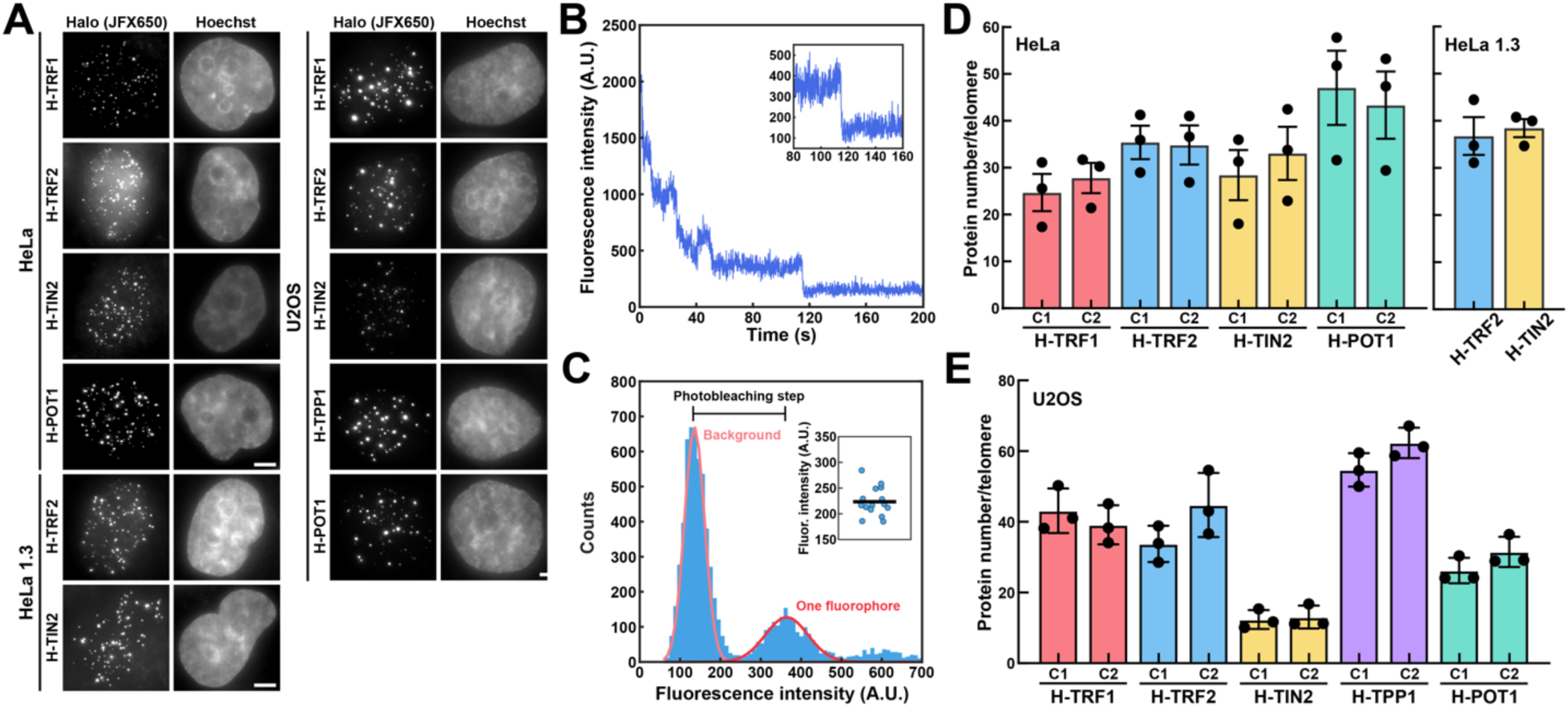
Shelterin protein abundance at telomeres. **(A)** Representative images of Halo-tagged shelterin proteins in HeLa, HeLa 1.3 and U2OS cells labeled with JFX650 Halo-ligand. Scale bar = 5 µm. **(B)** Photobleaching fluorescence time trace of representative Halo-TRF1 foci demonstrating single fluorophore photobleaching step. **(C)** Analysis of single fluorophore photobleaching step by fitting gaussian curves to the histogram representation of photobleaching function. Inset shows the analysis of multiple photobleaching functions to derive precisely the single fluorophore photobleaching step. **(D)** Quantification of telomeric protein number in HeLa and HeLa 1.3 cells based on dividing telomeric signal intensity by the intensity value of a single fluorophore (*N* = 3 biological replicates, median ± SD, see Figure S3E,F for distribution from a representative experimental replicate). **(E)** Quantification of telomeric protein number in U2OS cells based on dividing telomeric signal intensity by the intensity value of a single fluorophore (*N* = 3 biological replicates, median ± SD, see Figure S3G for distribution from a representative experimental replicate).

In total, these observations suggest that in HeLa cells the shelterin proteins are present in equal numbers with approximately 30 copies per telomeres. In U2OS cells, TIN2 appears to be present at substantially lower copy number than the other shelterin proteins. This might reflect that Halo-TIN2 is expressed at lower levels than endogenous TIN2 leading us to underestimate the number of TIN2 molecules. Alternatively, these results suggest that TPP1 and POT1 are recruited to telomeres via non-canonical mechanism in U2OS cells, for example by direct binding to single-stranded telomeric DNA.

### TRF2 displays faster dynamics at telomeres compared to the rest of the shelterin factors

To analyze the biophysical and biochemical properties of the shelterin proteins in living cells, we performed single-molecule live-cell imaging experiments using HILO (Highly Inclined and Laminated Optical sheet) microscopy (53) in combination with sparse protein labeling. This approach allowed us to localize and track individual HaloTagged shelterin factors over time in living cells (Fig. 4A,B, Movies S2-13). In all single-molecule imaging movies we observed mobile and static particles (Movies S3-13). To confirm that the static shelterin molecules are associated with telomeres, we first sparsely labeled the HaloTag with JFX650 Halo-ligand followed by quantitative labeling with the spectrally distinct JF549 HaloTag-ligand, to visualize telomeric foci (Fig. 4C). This analysis revealed that almost all static shelterin molecules were in close proximity to telomeric foci, consistent with their association with telomeres, rather than other unknown nuclear structures (Fig. 4D, Movies S14-17). For the Single-Particle Tracking (SPT) analysis, we used two different methods: The Spot-On algorithm (54) based on the evaluation of MSD (Mean-Square Displacement) distributions or a recently developed Bayesian approach using HMM (Hidden Markov Models) called ExTrack (55). Both of these methodologies allowed us to determine the fraction of immobile molecules, presumably representing molecules associated with telomeres or other chromatin loci, and the diffusion coefficients of mobile and static molecules, which provides a measure of the movement rate of the tagged proteins (Fig. 4E). For the analysis methods used in this study the number of molecular states has to be pre-defined. Using the Spot-On tool to fit the cumulative distribution function of the single-particle displacements the raw data fit well to a two-state model assuming a freely diffusing and telomere bound state (Fig. S4A). The same fit also closely matched the probability density function of the jump sizes (Fig. S4B,C), indicating that a two-state model is sufficient to describe the experimentally determined step size distributions. In U2OS cells, Spot-On analysis revealed that the majority (>60%) of TRF1, TIN2, TPP1, and POT1 were immobile and thus likely bound to telomeres (Fig. 4F). In contrast, TRF2 had a significantly lower fraction of static molecules (<30%), and most TRF2 molecules freely diffused through the nucleus (Fig. 4F). In HeLa cells the fraction of immobile shelterin molecules was lower than in U2OS cells for all proteins (Fig. 4G), consistent with a lower number of telomeric binding sites due to the shorter telomeres in HeLa cells, while protein levels were largely comparable (Fig. 2C,D). Importantly, the static fraction of TRF2 molecules was also significantly lower than the other shelterin proteins in HeLa cells (Fig. 4G). In addition, Halo-TRF2 also had a lower fraction of static molecules compared to Halo-TIN2 in HeLa 1.3 cells (Fig. S4F). Analysis of the diffusion rates revealed that static TRF1, TIN2, TPP1, and POT1 moved with comparable diffusion coefficients (D_bound_), while TRF2 was more mobile when bound to telomeres (Fig. 4H,I, Fig. S4F). Notably, the diffusion coefficient of freely diffusing particles (D_free_) was similar for all shelterin proteins in all cell lines analyzed (Fig. S4D-F). Since subunits of a multi-protein complex are expected to move with similar diffusion coefficients, these observations suggest that TRF1, TIN2, TPP1, and POT1 form a complex at telomeres, while the increased mobility of TRF2 indicates that it is not associated with the remaining shelterin subunits when bound to chromosome ends. To confirm this observation and to assure that the distinct biophysical properties of TRF2 are not a result of fusing it to the HaloTag, we integrated the HaloTag at the N-terminus of the *RAP1* locus in U2OS cells (Fig. S4G-I). In addition, we knocked-out RAP1 in cells expressing Halo-TRF2 in U2OS and HeLa backgrounds (Fig. S4H,I). The total cellular abundance of Halo-RAP1 (∼45,000 proteins per cell) was similar to that of Halo-TRF2 in U2OS cells (Fig. S4J). Since RAP1 recruitment to telomeres depends on its association with TRF2, we expected RAP1 to have similar diffusion properties to TRF2. The fraction of immobile RAP1 particles and the diffusion coefficient (D_bound_) of these molecules were indistinguishable from TRF2 and significantly different from TRF1 (Fig. 4J,K, Fig. S4K, Movie S18), consistent with TRF2 forming a constitutive complex with RAP1, and confirming that the diffusion dynamics of TRF2 at telomeres are distinct from TRF1, TIN2, POT1, and TPP1. In addition, the knock-out of RAP1 did not impact the diffusion properties of TRF2 (Fig. 4J,K, Fig. S4K, Movie S19). Importantly, the analysis of all of the data using ExTrack confirmed both the reduced fraction of immobile molecules and the increased telomeric diffusion coefficient of TRF2 compared to the other shelterin components (Fig. S4L,M). In total, these experiments suggest that at telomeres the shelterin complex largely exists in two distinct subcomplexes with different dynamic properties composed of TRF1-TIN2-TPP1-POT1 and TRF2-RAP1.

**Figure 4.**
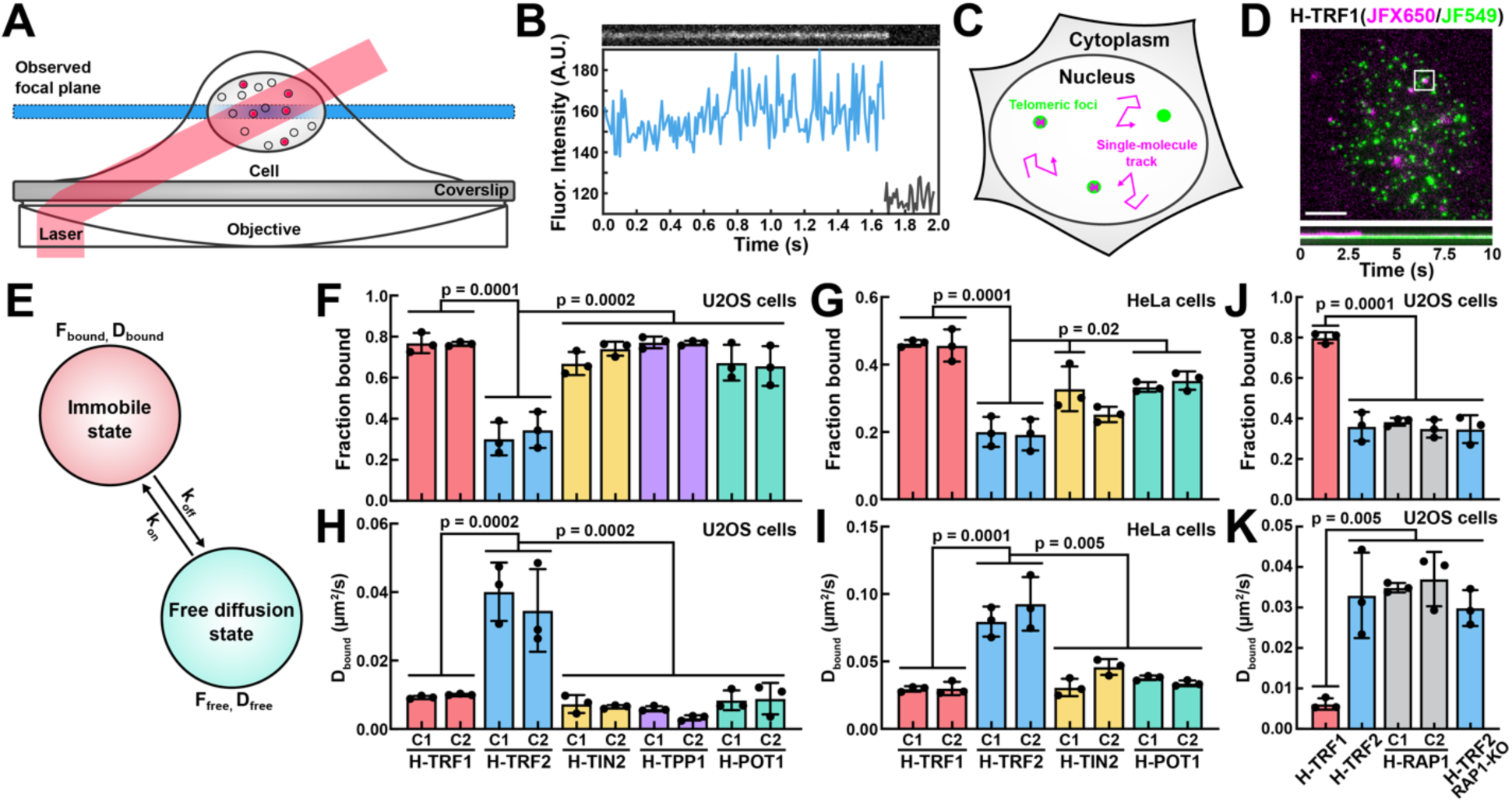
Single-molecule analysis of shelterin protein dynamics. **(A)** Illustration of HILO (Highly Inclined and Laminated Optical sheet) microscopy together with sparse labeling used for single-molecule live-cell imaging. **(B)** Raw intensity values of a representative single-molecule trajectory for Halo-TRF1 in U2OS cells monitoring statically associated molecule with telomere. **(C)** Visualization of combining sparse labeling for single-molecule detection (magenta) with dense labeling to mark telomeres (green). **(D)** Representative image (top) of HeLa cell expressing Halo-TRF1 sparsely labeled with JFX650 Halo-ligand (magenta) followed by dense labeling with JF549 Halo-ligand (green, Scale bar = 5 µm). Kymograph (bottom) of the telomere in the white box. **(E)** Illustration of the two-state model and the parameters associated with it used to analyze single-particle tracking experiments. **(F-G)** Quantification of the static fraction of HaloTagged shelterin components in **(F)** U2OS cells and **(G)** HeLa cells derived from a two-state fit of the cumulative distribution function of the single-particle displacements using the Spot-On software (*N* = 3 biological replicates, at least 30 cells per biological replicate, mean ± SD, one way ANOVA with Tukey’s post-hoc test). **(H-I)** Quantification of the diffusion coefficient of static HaloTagged shelterin components in **(H)** U2OS cells and **(I)** HeLa cells derived from a two-state fit of the cumulative distribution function of the single-particle displacements using the Spot-On software (*N* = 3 biological replicates, at least 30 cells per biological replicate, mean ± SD, one way ANOVA with Tukey’s post-hoc test). **(J-K)** Quantification of the **(J)** static fraction and **(K)** diffusion coefficient of static molecules of Halo-TRF1, Halo-TRF2, Halo-RAP1, and Halo-TRF2 in RAP1 knock-out U2OS cells derived from a two-state fit of the cumulative distribution function of the single-particle displacements using the Spot-On software (*N* = 3 biological replicates, at least 30 cells per biological replicate, mean ± SD, one way ANOVA with Tukey’s post-hoc test).

### TRF2 has a lower affinity for telomeres than TRF1 and the remaining shelterin subunits

To further analyze the biochemical properties of the shelterin components we determined the residence time of the HaloTagged shelterin proteins at telomeres using single-molecule imaging in living cells. The HaloTagged shelterin proteins were sparsely labeled with the highly photostable JF657 HaloTag-ligand, and imaged using a low laser power, 100 milliseconds exposure time, and in time intervals of 0.5 or 1.5 seconds to minimize photobleaching (Fig. 5A). The residence time of static shelterin proteins was determined using the distribution of trajectory lengths from single-particle tracking (Fig. 5B,C, Movie S20). The survival distribution of static shelterin protein dwell times fit well to two exponential decay functions (Fig. S5A), which could reflect shelterin proteins bound to telomeres and non-telomeric chromatin locations. Alternatively, the two binding times could reflect distinct telomeric binding sites from which the shelterin protein dissociate with different rates. In U2OS cells the time constant (which reflects the average association time) of long-lived binding events was lower for Halo-TRF2 (62 seconds), Halo-RAP1 (57 seconds), and Halo-TRF2 in RAP1 knock-out (54 seconds) cells compared to Halo-TRF1 (93 seconds), Halo-TIN2 (91 seconds), Halo-TPP1 (100 seconds), and Halo-POT1 (104 seconds) (Fig. 5D). In HeLa cells the time constant for long-lived binding events was comparable for all HaloTagged shelterin proteins (∼60 seconds, Fig. 5E). Importantly, the survival distributions generated from imaging data acquired in 0.5 s time intervals closely matched the data generated in 1.5 s intervals, demonstrating that photobleaching is not affecting this analysis (Fig. S5B,C). As a second approach to assess the affinity of the HaloTagged shelterin proteins for telomeric DNA we used ExTrack to determine the transition rate from the bound to the freely diffusing state derived from our fast single-molecule imaging and corresponding single-particle tracking analysis of shelterin protein dynamics (Fig. 4). The transition rate from the bound to the free state of Halo-TRF2 compared to the other shelterin components was 4-5 fold and 2-3 fold higher in U2OS and HeLa cells, respectively (Fig. 5F,G). The more pronounced difference in dissociation rates between Halo-TRF2 and other HaloTagged shelterin components shown by ExTrack compared to the residence time analysis is likely due to the fast image acquisition rate (100 frames per second) used for ExTrack that allowed us to detect extremely short-lived interactions which are not captured at low image acquisition rates. In total, these experiments suggest that TRF2 has a lower affinity for telomeric DNA than the other shelterin proteins analyzed, providing further evidence that shelterin forms two distinct subcomplexes at telomeres – TRF1-TIN2-TPP1-POT1 and TRF2-RAP1.

**Figure 5.**
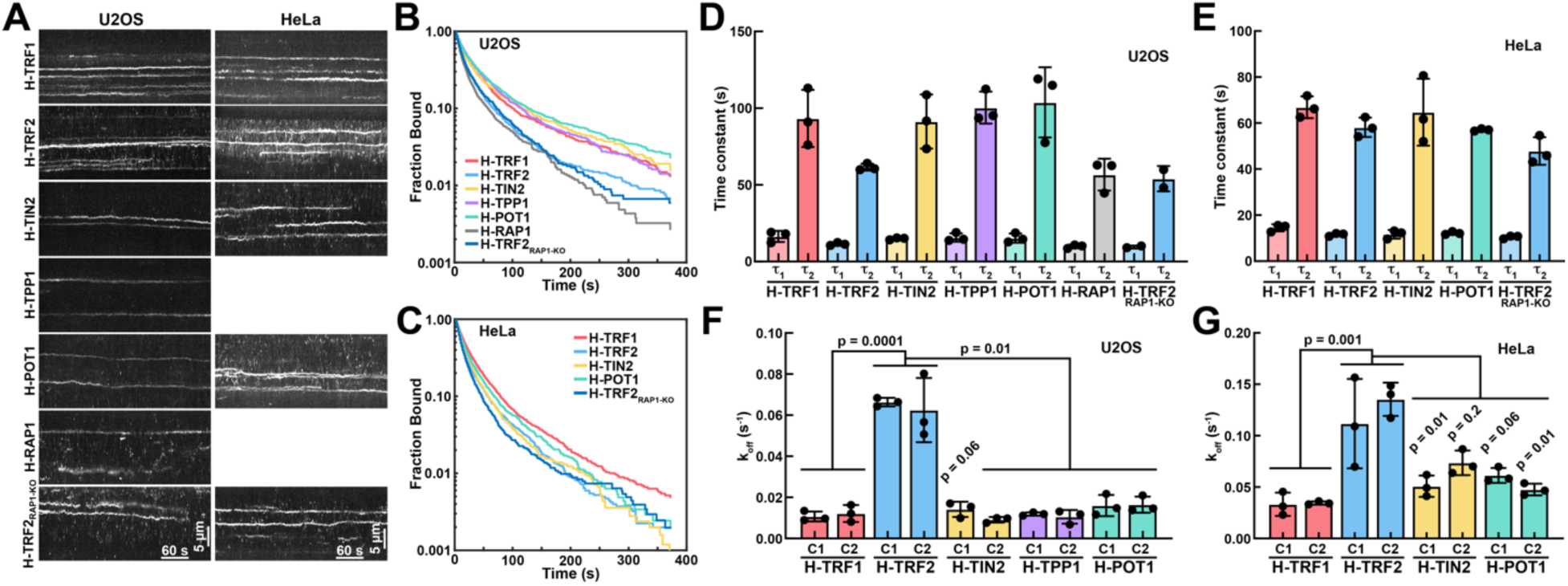
Single-molecule analysis of shelterin residence times at telomeres. **(A)** Kymographs of representative movies of the HaloTagged shelterin proteins labeled with JF657 HaloTag-ligand acquired at 1.5 s intervals. **(B-C)** Survival distributions of static shelterin molecules in **(B)** U2OS cells and **(C)** HeLa cells generated by single-particle tracking of immobile single-molecule signals of movies acquired in 1.5 s time intervals. **(D-E)** Time constants of short lived (τ_1_) and long lived (τ_2_) binding events derived from fitting the shelterin protein dwell time survival distributions from (D) U2OS cells and **(E)** HeLa cells using two exponential decay functions (*N* = 3 biological replicates, at least 10 cells per biological replicate, mean ± SD). **(F-G)** Transition rates of HaloTagged shelterin components from the bound to the freely diffusing state in **(F)** U2OS cells and **(G)** HeLa cells generated using ExTrack from rapid single-particle tracking experiments (Fig. 4, *N* = 3 biological replicates, at least 30 cells per biological replicate, mean ± SD).

### TRF1 and TRF2 form distinct subcomplexes that occupy different telomeric binding sites

To further test the model of two distinct shelterin subcomplexes at telomeres we set out to confirm the recruitment hierarchy of the shelterin proteins. Previous work by Frescas et al. demonstrated that TIN2 is largely recruited by TRF1 (21), which is consistent with our model that TRF1 and TRF2 form distinct complexes at telomeres. To define the recruitment hierarchy of the shelterin components we used a Halo-PROTAC ligand, which once it reacts with the HaloTag rapidly targets the HaloTag and its fusion partner for proteasomal degradation in living cells (Fig. 6A) (56). To analyze the telomere recruitment of the other shelterin proteins, we transiently expressed the mNeonGreen fusions of the shelterin factors in the cells treated with the Halo-PROTAC ligand. As expected, after Halo-TRF1 degradation TIN2, and POT1 were displaced from telomeres, while TRF2 recruitment was unaffected (Fig. 6B). In addition, RAP1 levels were reduced after TRF1 degradation (Fig. 6B). In contrast, TRF2 degradation had no visible effect on TRF1, TIN2, or POT1 recruitment (Fig. 6C). Only RAP1 localization to telomeres was diminished by TRF2 degradation, consistent with TRF2-RAP1 forming one complex (Fig. 6C). Similar to TRF1 depletion, degradation of TIN2 also eliminated POT1 recruitment and reduced RAP1 localization to telomeres (Fig. S6). This suggests that TRF2 over-expression can overcome the contribution of TIN2 to its recruitment to telomeres. However, RAP1 recruitment, even after over-expression, is partially dependent on TIN2, which suggests that TIN2 contributes to the recruitment of TRF2-RAP1 when TRF2 is expressed at endogenous levels. Interestingly, we also observed that over-expression of mNeonGreen-TRF1 was able to displace Halo-TRF1 from telomeres in control cells not treated with the Halo-PROTAC ligand (Fig. 6D). Similarly, expression of mNeonGreen-TRF2 removed Halo-TRF2 from telomeres (Fig. 6E). However, over-expression of mNeonGreen-TRF1 did not displace Halo-TRF2 from telomeres and mNeonGreen-TRF1 expression did not affect telomere occupancy of Halo-TRF2 (Fig. 6B,C). These observations are not only consistent with TRF1-TIN2-TPP1-POT1 and TRF2-RAP1 forming distinct sub-complexes, but they also suggest that TRF1 and TRF2 occupy distinct binding sites on telomeric chromatin.

**Figure 6.**
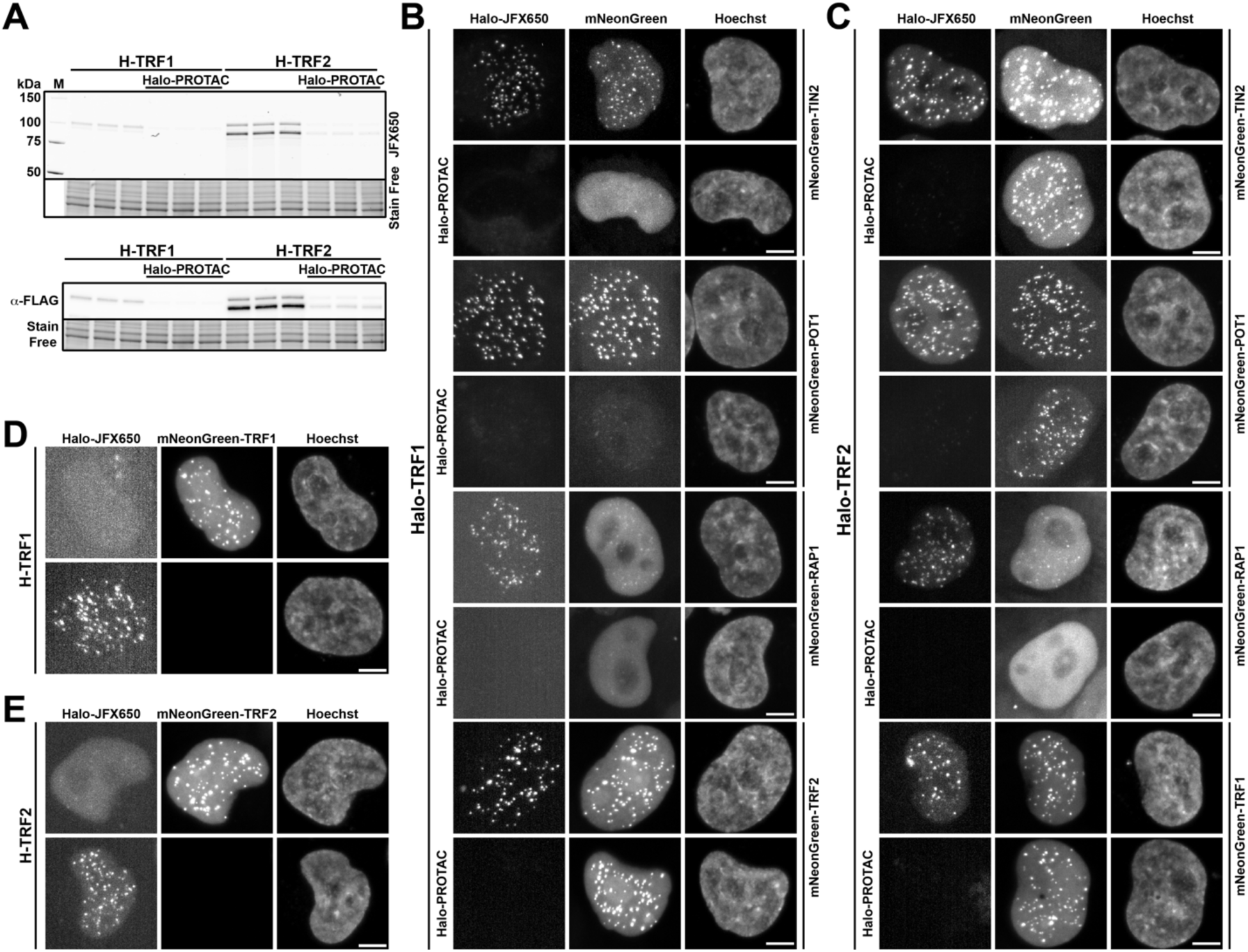
Analysis of the recruitment hierarchy of shelterin proteins to telomeres. **(A)** Fluorescence imaging (top) and anti-FLAG western blot (bottom) of the same gel loaded with cell lysates from HeLa cells expressing Halo-TRF1 or Halo-TRF2, untreated and treated with Halo-PROTAC for 18 hours. Cells were labeled with JFX650 HaloTag-ligand immediately prior to gel analysis. **(B)** Representative images of Halo-TRF1 HeLa cells transiently expressing mNeonGreen fusion proteins of TIN2, POT1, RAP1 and TRF2, untreated or treated with Halo-PROTAC for 18 hours. Cells were labeled with JFX650 HaloTag-ligand immediately before live-cell imaging. Scale bar = 5 µm. **(C)** Representative images of Halo-TRF2 HeLa cells transiently expressing mNeonGreen fusion proteins of TIN2, POT1, RAP1 and TRF1, untreated or treated with Halo-PROTAC for 18 hours. Cells were labeled with JFX650 HaloTag-ligand immediately before live-cell imaging. Scale bar = 5 µm. **(D)** Representative images of HeLa cells expressing Halo-TRF1 labeled with JFX650 HaloTag-ligand either untransfected (bottom) or transfected with a construct expressing mNeonGreen-TRF1. Scale bar = 5 µm. **(E)** Representative images of HeLa cells expressing Halo-TRF2 labeled with JFX650 HaloTag-ligand either untransfected (bottom) or transfected with a construct expressing mNeonGreen-TRF2. Scale bar = 5 µm.

## DISCUSSION

The data presented in this work completely re-envisions the architecture of telomeres in human cancer cells. Using a panel of genome edited cell lines expressing HaloTagged shelterin components from their endogenous loci in three different cancer cell lines, we precisely determined the abundance of the shelterin proteins at chromosome ends, defined their diffusion dynamics at telomeres, and demonstrated that TRF1 and TRF2 occupy distinct binding sites even though they both bind to telomeric repeat DNA *in vitro*. Moreover, the HaloTagged shelterin cell lines generated in this work will serve as a valuable resource for the scientific community to facilitate future studies of telomere function.

### Overall architecture of telomeres in cancer cells

To define the molecular architecture of telomeres in cancer cells we determined the overall cellular abundance of the shelterin proteins and their copy number at telomeres. Our experiments demonstrate that TRF1 is present at ∼10-12,000 molecules per cell in both U2OS and HeLa cells, while TRF2 expression levels are at least 5-fold higher (∼60,000 and ∼85,000 proteins per cells for U2OS and HeLa cells, respectively). In HeLa cells TIN2 and POT1 were present at similar levels (∼35,000 proteins per cells) and in U2OS cells TIN2, POT1, and TPP1 protein levels were also comparable (∼10-20,000 proteins per cells). For TRF1, TRF2, TPP1, and POT1 these protein levels align well with previous observations made by Takai et al (45). However, the levels of TIN2 in U2OS cells determined here are around 10-fold lower than previously suggested (∼10,000 compared to 125,000 proteins per cell) (45). It is important to note that we were unable to confirm that Halo-TIN2 is expressed at similar levels as endogenous TIN2. It is therefore possible that Halo-TIN2 is expressed at lower levels than untagged TIN2. Moreover, TIN2 levels were not previously determined in cells using ALT for telomere maintenance. Our Halo-TIN2 cell lines did not have a growth defect, and we did not detect telomere dysfunction induced foci, which demonstrates that Halo-TIN2 is fully functional in telomere end protection.

Our fluorescence photobleaching approach allowed us to determine the copy number of the shelterin proteins directly at telomeres. Our observations demonstrate that the median number of shelterin proteins in HeLa cells is ∼30 copies per telomere. Considering a telomere length of ∼4,000 nucleotides (57) and a spacing of telomeric nucleosomes of 157 nucleotides (i.e. 27 nucleosomes per telomere) (58), these observations suggest that telomeres are bound by one TRF1 and TRF2 dimer for every two telomeric nucleosomes (Fig. 7). Even though U2OS and HeLa 1.3 cells have 4 to 5-fold longer telomeres than the HeLa cells used in this study, the abundance of TRF1 and TFR2 is only slightly higher in these cell lines (∼40 copies per telomere). This suggests that the shelterin proteins are likely more spread out on chromosome ends in cells with long telomeres. A lower shelterin density on long telomeres could have important implications for telomerase recruitment, since association with an internally bound TPP1 molecule could make it more difficult for telomerase to localize to the chromosome end for elongation.

**Figure 7.**
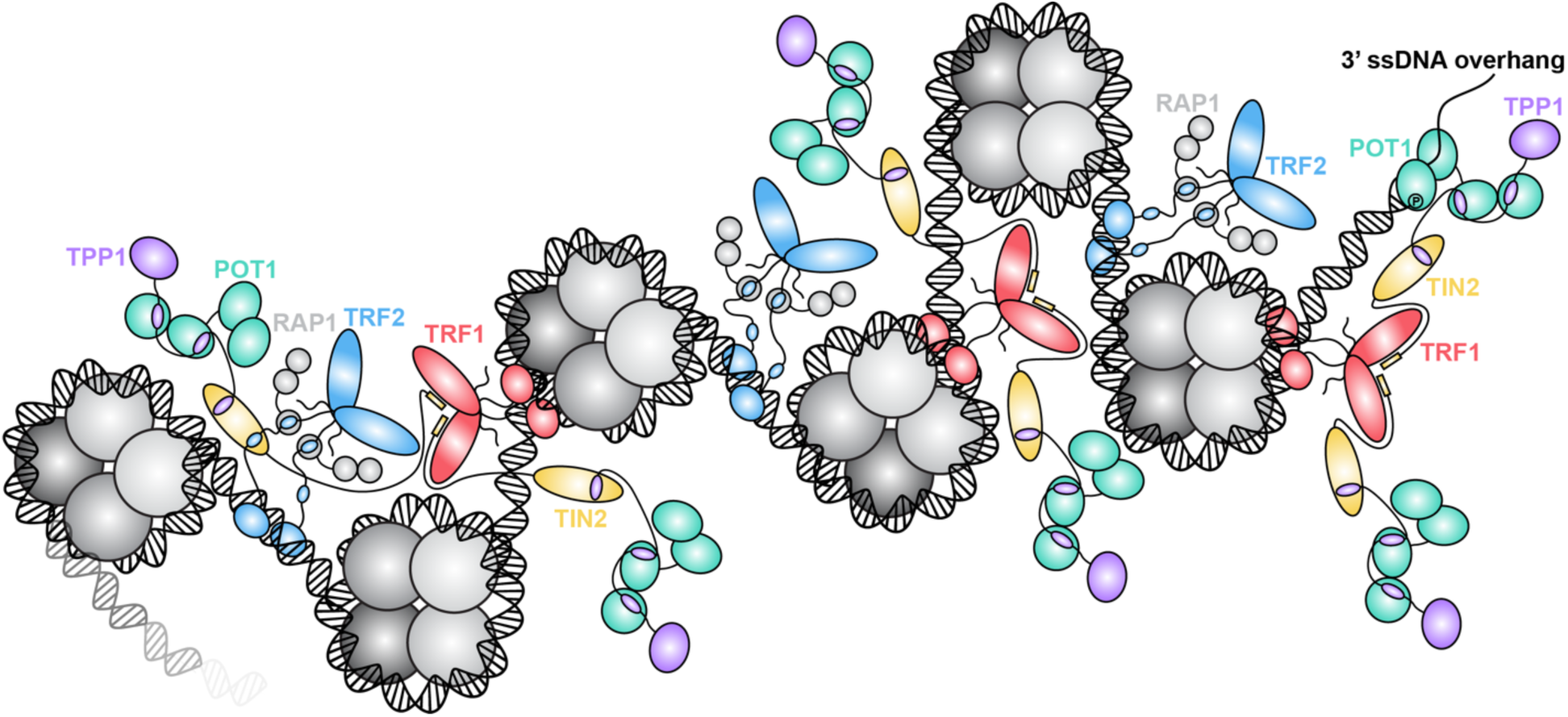
Model of telomere architecture in HeLa cells with short telomeres. The shelterin complex associate with telomeres predominantly in a form of two distinct subcomplexes: TRF1-TIN2-TPP1-POT1 binding the entry/exit sites of nucleosomes, and TRF2-RAP1 binding likely the linker DNA between nucleosomes. TRF2 can transiently associate with TIN2 to stabilize its interaction with the chromosome end. Both subcomplexes are dimeric and are presented at telomeres in equal numbers binding approximately every other nucleosome.

In U2OS cells we found that the median number of telomere-associated Halo-TIN2 molecules was 10. This number is substantially lower than the number of TPP1 and POT1 molecules, which were present at approx. 60 and 30 proteins per telomere in U2OS cells, respectively. As outlined above, it is possible that we are underestimating the number of TIN2 molecules at telomeres because Halo-TIN2 might be expressed at lower levels than endogenous TIN2. Alternatively, it is possible that in ALT cells the TPP1-POT1 complex can be recruited to or retained at telomeres via its interaction with single-stranded stretches of telomeric DNA without maintaining an association with TIN2. Therefore, future work is needed to further define the shelterin stoichiometry and its function in ALT cells.

The telomeric copy number of the shelterin proteins determined here are up to 50-fold lower than previously reported numbers (ranging from 65 for POT1 to 720 copies per telomere for TIN2), calculated by measuring the absolute cellular abundance and chromatin associated fractions of shelterin proteins (45). It is possible that the presence of the HaloTag reduces the abundance of shelterin proteins at telomeres. However, it is unlikely that all shelterin proteins would be impacted in a similar way by the fusion to the HaloTag. Our live-cell single-molecule imaging experiments indicate that a substantial nuclear fraction (60-80% in HeLa cells) of the shelterin proteins are not associated with chromatin. It is therefore possible that determining the chromatin bound fraction of the shelterin proteins using cell lysis and fractionation overestimated the percentage of the shelterin subunits associated with telomeres. Calculation of the telomeric abundance of TRF1 in HeLa cells using the total cellular abundance and chromatin bound fraction determined in this study (∼10,000 molecules per cells * 46% fraction bound / 142 telomeres) results in 33 TRF1 molecules per telomere, which closely matches our photobleaching results and increases our confidence in our findings.

### TRF1 and TRF2 form subcomplexes with distinct telomeric binding sites

A large number of previous studies has addressed the biochemical basis of shelterin complex formation. When all subunits (TRF1, TRF2, RAP1, TIN2, TPP1, POT1) are combined, shelterin can form a fully dimeric complex (41). Similarly, TRF1, TIN2, and TPP1 were shown to form a complex with 2:2:2 stoichiometry (44). In contrast, TRF2-RAP1 can form a 2:1 complex with the remaining shelterin subunits, containing a dimer of TRF2-RAP1 and a single copy of TIN2, TPP1, and POT1 (43). In cells, the work presented here and previous work by Frescas et al. demonstrates that TRF1 is responsible for the recruitment of TIN2 to telomeres, and TRF2 is not required for TIN2 localization to chromosome ends (21). Therefore, TIN2 must be recruited to telomeres via its interaction with the TRFH domain of TRF1. This also suggests that the TRFH domain of TRF2 rarely if ever interacts with TIN2 at telomeres in cells. The TRFH domain of TRF2 is therefore likely always available to associate with other co-factors, for example Apollo or SLX4, to rapidly and dynamically recruit critical chromosome end protection proteins to telomeres (13,14,16,17).

In HeLa cells the number of telomeric TIN2 and POT1 molecules closely matched the number of TRF1 and TRF2 molecules. This observation is in principle consistent with the formation of a fully dimeric complex composed of two copies of TRF1, TRF2, RAP1, TIN2, TPP1, and POT1. However, our single-molecule imaging experiments demonstrate that TRF2 and RAP1 are substantially more dynamic when bound to telomeres than TRF1, TIN2, TPP1, and POT1, which suggests that they form two distinct subcomplexes. This is further corroborated by the higher dissociation rate constant of TRF2 from telomeres compared to TRF1-TIN2-POT1-TPP1, and the lower residence time of TRF2 at telomeres in U2OS cells. Altogether our observations suggest that TRF1, TIN2, TPP1, and POT1 form a dimeric complex containing two copies of each subunit, which tightly associates with telomeric DNA. In contrast, the TRF2-RAP1 complex is more dynamically bound to telomeres, potentially rapidly repositioning between different telomeric binding sites. Importantly, the work presented here, and previous work by others, demonstrates that TIN2 contributes to the recruitment of TRF2-RAP1 to telomeres (22,23,34,59), which must be mediated by the interaction formed between the hinge region of TRF2 and the N-terminal TRFH domain of TIN2. Consistent with a highly dynamic interaction, the affinity of the TIN2’s TRFH domain for the TRF2’s hinge region is in the low micromolar range (18), which is substantially less stable than the interaction between TIN2 and the TRFH domain of TRF1 (15). TRF2-RAP1 could therefore rapidly reposition on telomeric DNA by forming transient interactions with both telomeric DNA, mediated by the MYB domain of TRF2, and the TRFH domain of TIN2 (Fig. 7). This would allow TRF2 to rapidly scan along telomeric DNA to drive t-loop formation (25–28,60), localize Apollo to the chromosome end to facilitate end resection (15,16,61), or to prevent the recruitment of the NHEJ machinery to blunt ended telomeres (4,62,63).

The final piece of evidence that TRF1 and TRF2 form separate subcomplexes is that they occupy distinct telomeric binding sites in cells. Our experiments demonstrate that over-expression of TRF1 or TRF2 can displace itself from telomeres but is unable to outcompete the other shelterin component. This strongly suggests that TRF1 and TRF2 bind to distinct regions of telomeric chromatin. A possible explanation was recently suggested by structural analysis of telomeric nucleosomes bound to the MYB domain of TRF1 (44). The MYB domain of TRF1 was found to specifically associate with the entry and exit sites of telomeric nucleosomes, while TRF2 was not observed to bind in a similar manner (44,64). While this model provides a straightforward explanation for the inability of TRF2 to displace TRF1, it is less clear why TRF1 is unable to displace TRF2 from telomeres. If TRF2 associates with naked telomeric repeat DNA, for example the short linker DNA between nucleosomes, sufficiently high levels of TRF1 should be able to outcompete TRF2. One possibility is that binding to TIN2 could retain TRF2 at telomeres when TRF1 is over-expressed and fully occupies potential telomeric DNA binding sites of TRF2. Alternatively, TRF2 could specifically recognize other features of telomeric chromatin (e.g. bent DNA on the outside of nucleosomes) that TRF1 is unable to associate with. Further work is necessary to dissect the relative contribution of TIN2 and telomeric DNA binding to the recruitment of TRF2 to telomeres.

In total our results demonstrate that telomeric chromatin in HeLa cells is densely covered with shelterin proteins. The TRF1-TIN2-TPP1-POT1 complex is tightly associated with telomeric chromatin to facilitate telomere replication, prevent ATR activation, and to recruit telomerase to telomeres. In contrast, the TRF2-RAP1 complex dynamically associates with telomeres to allow it to recruit a variety of co-factors to chromosome ends that prevent ATM activation and to inhibit NHEJ. In ALT cells, chromosome ends are more sparsely covered by the shelterin complex and stoichiometry of the shelterin components is distinct from telomerase positive cancer cells.

## ACKNOWLEDGMENTS

This work was supported by grants from the NIH (R01 GM141354, DP2 GM142307) to J.C.S. We thank to Luke Lavis lab (HHMI Janelia Research Campus) for providing Janelia Fluor ligands. We kindly thank Jan Karlseder for providing the TRF1 antibody used in this study. All flow cytometry data were obtained using instrumentation in the MSU Flow Cytometry Core Facility and we especially thank to Matthew Bernard for providing service regarding cell sorting and flow cytometry analysis. The flow cytometry facility is funded in part through the financial support of Michigan State University’s Office of Research & Innovation and Colleges of Osteopathic Medicine, Human Medicine, Veterinary Medicine, Natural Sciences, and Engineering. We thank to MSU IQ microscopy core facility for supporting our work.

## AUTHOR CONTRIBUTIONS

Conceptualization: T.J. and J.C.S.; Experiments: T.J. and G.I.P.; Data Analysis: T.J.; Writing: Original Draft: T.J.; Writing: Review and Editing: T.J. and J.C.S.

## COMPETING INTERESTS

The authors declare no competing financial interests.

## DATA AVAILABILITY

All primary data will be provided upon request by J.C.S.

## CODE AVAILABILITY

All code used in this study has been deposited on GITHUB (https://github.com/SchmidtLabMSU/ShelterinAnalysis).

## MATERIALS AND METHODS

### Cell lines maintenance

U2OS cells were grown in RPMI 1640 media (Gibco) supplemented with 10% FBS (Gibco), 100 U ml^−1^ penicillin and 100 µg ml^−1^ streptomycin (Gibco) at 37°C and 5% CO_2_. All generated HeLa cell lines were based on HeLa-EM2-11ht(47) and were grown in DMEM media (Gibco) supplemented with 10% FBS, 100 U ml^−1^ penicillin and 100 µg ml^−1^ streptomycin (Gibco) at 37°C and 5% CO2.

### Generation of HaloTagged cell lines and plasmid construction for genome editing

To insert HaloTag at the 5’ end of all shelterin genes, the CRISPR/Cas9 genome-editing strategy was used with following procedure. All sgRNAs were cloned into BpiI-digested pX330-U6-Chimeric-BB-CBh-hSpCas9 backbone (Addgene #42230, gift from Feng Zhang). Plasmids used as donors for homology-directed repair (HDR) were prepared using Gibson assembly (NEB) with HpaI-digested pFastBacDual backbone, HaloTag insert, left and right homology arms. Homology arms were either PCR amplified from genomic DNA (*TRF1, TRF2, TPP1, POT1* genes) or ordered as g-Blocks from IDT (*TIN2, RAP1* genes). HaloTag insert consisted of 3xFLAG-HaloTag sequence, followed by a short linker sequence including TEV cleavage site. Sanger sequencing was used to confirm the correct cloning of plasmids. For genome editing, cells were plated into 6-well plate to reach ∼70% confluency next day. Cells were transfected with a mixture of 1.25 µg of HDR donor plasmid, 1.25 µg of sgRNA plasmid and 7.5 µl of Lipofectamine 2000 (Invitrogen). After 48-72 h of transfection, cells were fluorescently labeled with Halo-ligand JFX650 and single-cell sorted into 96-well plates using FACS. Non-edited cells stained with JFX650 Halo-ligand were used as a negative control for acquiring background fluorescence. Single-cell clones were propagated into 24-well plates and screened for homozygous insertion by in-gel fluorescence and genomic PCR followed by Sanger sequencing. To knock-out RAP1 in Halo-TRF2 cell lines, cells were transfected (same protocol as above) with plasmid eSpCas9-ATP1A1-G2-Dual-sgRNA (Addgene #86612, gift from Yannick Doyon) which enabled for marker-free co-selection for NHEJ-based gene editing(65). The sgRNA targeting Exon 1 of *RAP1* gene was cloned into the BbsI-linearized eSpCas9 vector. Three days post transfection, cells were selected with 0.5 µM ouabain (Sigma-Aldrich) and then single-cell diluted into 96-well plates. Clones containing *RAP1* knock-out were screened and validated by western blotting. Cloning of the mNeonGreen fused shelterin proteins was performed by Gibson assembly into pRK2 backbone (original pHTN HaloTag CMV-neo, Promega #G7721).

### Western blotting

Cells grown at 24-well plate were lysed with Laemmli sample buffer (BioRad) and subjected to SDS-PAGE using 4-15% TGX Stain-Free polyacrylamide gels (BioRad). Resolved proteins were transferred to a nitrocellulose membrane using Trans-Blot Turbo system (BioRad). Membrane was blocked with 5% milk in PBS-T (0.05% Tween 20) for 1 hour at RT followed by incubation with primary antibody (diluted in 2.5% milk in PBS-T) O/N at 4°C. Then, membrane was washed 3 x 5 min in PBS-T and incubated for 1 hour at RT with secondary antibody anti-mouse (Invitrogen #31439) or anti-rabbit (Invitrogen #31460) conjugated with HRP diluted 1:4,000 in 5% milk in PBS-T. Finally, membrane was washed 3 x 10 min in PBS-T and imaged for chemiluminescence using ChemiDoc system (Biorad) and SuperSignal West Femto Maximum Sensitivity Substrate (Thermo Scientific). Signal intensities were analyzed using ImageQuant software (Cytiva). Primary antibodies used in this study: anti-TERF2 (Novus #NB100-56506, 1:1,500 dilution), anti-TERF2IP (Novus #NB100-292, 1:1,000 dilution), anti-TPP1 (Bethyl #A303-069A, 1:2,000 dilution), anti-POT1 (Proteintech #10581-1-AP, 1:750 dilution), anti-TRF1 (raised in rabbit, 1:1,000 dilution, kind gift from Jan Karlseder), anti-FLAG conjugated with HRP (Sigma-Aldrich, #A8592, 1:2,000 dilution).

### Preparation of 6xHis-3xFLAG-HaloTag for in-gel protein abundance measurement

The plasmid carrying 6xHis-3xFLAG-HaloTag was kindly provided by Dr. Youmans and Dr. Cech (University of Colorado Boulder, Boulder, CO, USA). The construct was transformed into BL21-Star (DE3) *E. coli* (Invitrogen) by heat-shock protocol. Cells were grown in Luria-Bertani media containing 100 µg ml^−1^ of ampicillin at 37°C and 200 rpm until O.D._600_ = 0.8, then temperature was lowered to 18.5°C and the expression was induced with 0.5 mM IPTG. Cells were cultured O/N and harvested next morning by centrifugation (4,000 g x 10 min) at RT and pellet was stored at -80°C. All purification steps were carried out at 4°C unless otherwise stated. The bacterial pellet (12 g) was dissolved in 80 ml of lysis buffer (50 mM sodium phosphate pH 8, 500 mM NaCl, 10 mM Imidazole, 5% glycerol, 0.5% Tween 20, 1 mM DTT, 0.5 mM PMSF, 0.5 mg ml^−1^ lysozyme) and sonicated for 3 min (process time) with 40% amplitude, 1 s pulse ON and 3 s pulse OFF (Fisherbrand Model 505, 0.5-inch tip). The suspension was cleared by centrifugation (19,000 g x 1 h). The supernatant was loaded on a gravity column containing 2.5 ml of Ni Sepharose 6 Fast Flow resin (Cytiva) and incubated for 30 min. Then, column was washed with 3 CV of lysis buffer and protein was eluted with elution buffer (50 mM sodium phosphate pH 7.5, 500 mM NaCl, 300 mM Imidazole, 5% glycerol). Sample was concentrated to 5 ml using Vivaspin 20 10 kDa MWCO (Cytiva), centrifuged (4,000 g x 10 min) and loaded onto HiLoad 16/600 Superdex 75 pg column (GE Healthcare Life Sciences) equilibrated in FPLC buffer (50 mM sodium phosphate pH 7.5, 150 mM NaCl, 5% glycerol). Flow-rate for size-exclusion chromatography was set to 1 ml min^-1^. Peak fractions according to A_280_ were analyzed by SDS-PAGE, pooled and concentrated to 13.5 mg ml^−1^ using Vivaspin 20 10 kDa MWCO (Cytiva). Final sample was aliquoted, flash-frozen in LN_2_ and stored at -80°C. To fluorescently label the prepared protein with JF657 Halo-ligand, the protein was diluted with FPLC buffer to 3 mg ml^−1^ and mixed with 3-fold molar excess of JF657 Halo-ligand (resuspended in 1/10 of a protein volume of FPLC buffer supplemented with 10% DMSO). Mixture was incubated in dark for 3 hours at 4°C while slowly rotating. Then, the mixture was centrifuged (15,000 g x 10 min) at 4°C and purified by size-exclusion chromatography (same protocol as before) to remove the excess dye. Peak fractions according to A_280_ were pooled and concentrated using Vivaspin 6 10 kDa MWCO (Cytiva) to 10 µM. The degree of labeling was ∼91% (determined by UV-Vis spectroscopy). The extinction coefficients used for calculations were ε_280_ = 67,380 M^−1^ cm^−1^ for the 6xHis-3xFLAG-HaloTag (analyzed using Expasy ProtParam tool (66)) and ε_max_ = 137,000 M^−1^ cm^−1^ for the JF657 Halo-ligand (67). The correction factor for JF657 equaled to 0.0267, which was experimentally measured as A_280_/A_max_ ratio of the ligand. Fluorescently labeled protein was diluted to 10 nM according to A_max_ and aliquoted by mixing 10 µl of protein + 10 µl of 2x Laemmli sample buffer (Biorad). Therefore, each aliquot represented 100 fmol of fluorescently labeled Halo standard. Aliquots were boiled for 10 min at 95°C, flash-frozen in LN_2_ and stored at -80°C until use.

### In-gel fluorescence for quantification of protein total cellular abundance

Cells (100,000 for U2OS and 150,000 for HeLa) were plated into 24-well plate. Next day, cells were labeled with 250 nM JF657 Halo-ligand for 1 h, washed with PBS and incubated with a fresh media for additional 15 min. Then, cells were washed twice with PBS and lysed in 60 μl Laemmli sample buffer (BioRad) and boiled for 10 min at 95°C prior to SDS-PAGE. Prepared Halo standards (section above) were supplemented with a known amount of lysed parental cells (in Laemmli sample buffer), serially diluted, and loaded together with samples on 4-15% TGX Stain-Free polyacrylamide gel (BioRad). In-gel fluorescence was imaged using ChemiDoc system (Biorad) with Cy5.5 filter set, and total protein loading was imaged using Stain free filter with UV activation. To measure the protein abundance, the JF657 signal intensity was related to the standard curve obtained from serially diluted Halo standards. Similarly, the number of loaded cells were related to the standard curve based on Stain free signal of prepared standards. All quantifications were performed using ImageQuant software (Cytiva). For TEV correction, plated cells were labeled with JF657 Halo-ligand and washed as above. Then, cells were lysed in 50 μl RIPA buffer (w/o SDS). Samples were split half, supplemented with protease inhibitor cocktail (Sigma-Aldrich, #P8340) and one set of samples were treated with 5 units of TEV protease (NEB, #P8112) for 30 min on ice. After TEV cleavage, treated and untreated samples were mixed with 2x Laemmli sample buffer (Biorad), boiled for 10 min at 95°C and separated by SDS-PAGE. In-gel fluorescence was detected using the same procedure as above and TEV correction factors were obtained as a ratio between the signal intensity of uncleaved vs cleaved HaloTagged protein normalized to Stain free signal. All experiments were performed in triplicate.

### Flow-FISH for measuring relative telomere length

U2OS or HeLa cells were split by trypsinization and 3x10^6^ cells were transferred to a separate tube, washed with PBS and the cell pellet was resuspended in 1.2 ml of PBS. While slowly vortexing, 2.8 ml of 100% EtOH was added dropwise to fix cells (70% EtOH in final volume). Fixed cells were stored in the freezer until use. For the telomeric FISH staining, cells were washed with PBS and resuspended in 100 µg ml^−1^ of RNase A in PBS followed by incubation at 37°C for 20 min. Then, cells were washed with PBS and permeabilized in PBS + 0.05% Triton X-100 for 10 min at RT. After permeabilization, cells were washed with PBS and resuspended in preheated (80°C for 5 min) hybridization buffer (20 mM Tris pH 7.4, 60% formamide, 0.5% blocking solution (Roche #11096176001) and 1:300 dilution of PNA TelC-AF488 probe (PNA Bio #F1004)). Mixture was transferred to PCR tubes and incubated in thermocycler at 80°C for 10 min followed by 2 hour incubation at RT in dark. After the hybridization, samples were transferred to 96-well plate and washed 2 x 10 min with preheated (60°C) 2x SSC + 0.1% Tween 20 buffer. Then, cells were washed 2 x 5 min with PBS and finally resuspended in 150 µl of PBS and stored at 4°C covered in aluminum foil for the flow cytometry measurement next day. All centrifugation steps were done at 300 g x 3 min at RT with a bucket-swing rotor. Flow cytometry measurement was performed using spectral flow cytometer Cytec Aurora (Cytec). Samples were at 96-well plate. Acquired data were unmixed in SpectroFlo software (Cytec) and further analyzed in FCS Express (De Novo Software).

### Immunofluorescence and fluorescence in-situ hybridization for TIF analysis

Parental or generated HaloTagged cell lines were plated into 6-well plate (250,000 cells for U2OS and 400,000 cells for HeLa) containing sterilized 18 x 18 mm #1.5 glass coverslips (Corning). The next day, cells were labeled with 250 nM of JFX650 Halo-ligand for 30 min in complete media followed by two washes and a 15 min incubation with fresh complete media. After that, cells were washed with PBS and fixed with 4% formaldehyde in PBS for 10 min at RT. After fixation, cells were washed twice with PBS and permeabilized with 0.1% Triton X-100 in PBS for 10 min at RT. Then, cells were washed twice with PBS and blocked with 3% BSA + 0.05% Tween 20 in PBS for 45 min at RT followed by the incubation with anti-53BP1 antibody (Novus, #NB100-304) diluted 1:1,000 in blocking buffer for 1 hour at RT. Cells were washed 3 x 5 min with PBS-T (0.05% Tween 20) followed by the incubation with secondary anti-rabbit antibody conjugated with Cy3 (Invitrogen, #A10520) diluted 1:500 in blocking buffer for 1 hour at RT. Then, cells were washed 3 x 5 min with PBS-T and fixed again with 4% formaldehyde in PBS for 5 min at RT. After fixation, cells were dehydrated with increasing concentration of EtOH (70%, 85% and 100%) for 2 min each. Coverslips were then dried and transferred into preheated (80°C) glass slides and incubated with a preheated (80°C for 5 min) hybridization buffer (20 mM Tris pH 7.4, 60% formamide, 0.5% blocking solution (Roche #11096176001) and 1:200 dilution of PNA TelC-AF488 probe (PNA Bio #F1004)) for 10 min at 80°C followed by 2 hour incubation at RT in dark. After hybridization, cells were washed 2 x 10 min with preheated (60°C) 2x SSC + 0.1% Tween 20 buffer. Then, cells were incubated with Hoechst dye (1:10,000 dilution in 2x SSC) for 2 min at RT to stain nuclei. Finally, cells were washed with 1x SSC and MilliQ water, dried, and mounted onto glass slides with ProLong diamond antifade mountant (Invitrogen #P36970) followed by O/N curing at RT in dark. Samples were imaged the next day using 3i spinning-disc confocal microscope with SoRa modality. Acquired Z-stacks for all channels were processed in ImageJ to obtain maximum intensity projections, which were used for subsequent automatic high-throughput analysis using custom MATLAB script. Briefly, nuclei channel was segmented using Cellpose machine-learning algorithm (68) in MATLAB, telomeric and 53BP1 foci were localized using ComDet plugin (69) in ImageJ. Then, the number of TIFs per each segmented cell was determined as a number of colocalization events between 53BP1 and TelC. For each sample, minimum of 100 cells were included into the analysis. Statistical analyses were obtained using the Mann-Whitney test in R.

### Microscopy setup for live and fixed cell imaging

Two distinct microscopy systems were used for imaging either live or fixed cells. First, the Olympus IX83 inverted microscope equipped with cellVivo incubation system (temperature, humidity and CO_2_ controlled), four laser lines (405, 488, 561 and 640 nm), X-cite Turbo LED excitation source, Olympus UPlanApo-HR 60x/1.5 NA TIRF objective, Twin-cam beamsplitter (Cairn Research) and Hamamatsu Orca-Fusion BT sCMOS or Hamamatsu Orca-Quest qCMOS cameras. The second system was a 3i spinning-disc (Yokogawa) microscope with confocal SoRa modality equipped with incubation chamber (temperature, humidity and CO2 controlled), four laser lines (445, 488, 561 and 638 nm), Zeiss C PlanApo 63x/1.42 NA objective and Hamamatsu Orca-Fusion BT sCMOS camera. All live-cell imaging experiments were carried out at 37°C, 5% CO_2_ and 95% humidity-controlled environment to mimic the conditions used for cell culturing.

### Measuring the shelterin abundance at telomeres by fluorescence photobleaching

HaloTagged shelterin cell lines (25,000 U2OS and 45,000 HeLa cells) were plated into 24-well #1.5H glass bottom dish (Celvis #P24-1.5H-N). To synchronize cells at the onset of S-phase, cells were incubated the next day with 2 mM thymidine for ∼18 hours. Then, cells were washed twice with complete media and incubated for 9 hours to complete the DNA replication. Cells were then blocked again with 2 mM thymidine for ∼18 hours. During the last hour of incubation, cells were labeled with 250 nM Halo-JFX650 ligand for 45 min in the presence of 2 mM thymidine. After labeling, cells were washed twice and incubated for additional 15 min in a fresh complete media containing 2 mM thymidine. Then, cells were washed twice and fixed with 4% formaldehyde for 10 min at RT followed by two-time wash with PBS and nuclei staining for 4 min at RT using Hoechst dye (diluted 1:5,000 in PBS). Cells were washed twice with PBS and imaged immediately using the Olympus IX83 microscope (equipped with Orca-Quest camera). Z-stacks of Halo channel (51 planes) were imaged using 640 nm laser line under epi-illumination with these settings: 100% laser power, 40 ms exposure time (HeLa cells) or 65% laser power, 25 ms exposure time (U2OS and HeLa 1.3 cells). Subsequently, the Z-stack of Hoechst channel was acquired using LED 385 nm excitation light. After collecting Z-stacks for at least 60 cells per sample, the photobleaching time-lapse movies (continuous imaging of one plane capturing 8,000 frames for HeLa or 15,000 frames for U2OS) of Halo-TRF1 foci were acquired using the same exposure settings. Halo-TRF1 cells were chosen due to their specific telomeric shelterin foci formation with minimal nuclear background signal. The analysis for quantifying the number shelterin proteins bound to telomeres was done in two steps using custom written MATLAB scripts. Firstly, the photobleaching movies were used to determine the fluorescence intensity of a single fluorophore. At least 20 Halo-TRF1 foci were cropped and their maximum intensity profiles over time were saved using ImageJ. From these traces the background fluorescence (mean intensity over time of nucleoplasm ROI) was subtracted. Then, each photobleaching function was plotted as a histogram where the position of maximum count was detected and corresponding peak was fitted with gaussian distribution, which represented the fluorescence background level. Next, the closest peak to the background distribution was localized and fit with another gaussian function, which represented the fluorescence intensity distribution of a single fluorophore. The difference between the mean values of those gaussian distributions was assigned as a photobleaching step-size, which was used in the next step to extract the number of fluorophores within the foci from its initial foci intensity value. The second step consisted of maximum intensity projection of all acquired Z-stacks with subsequent foci localization using ComDet (69) plugin in ImageJ. Nuclei channel was segmented using Cellpose (68) machine-learning algorithm in MATLAB. Finally, the fluorescence intensity for each individual shelterin telomeric foci was background subtracted and divided by photobleaching step-size, which allowed to extract the number of HaloTagged shelterin molecules within a respective telomeric foci with single cell resolution in a high-throughput manner.

### Flow cytometry

To analyze the cell cycle profile of cells treated with double-thymidine block used for the analysis of shelterin abundance at telomeres, cells were plated in 6-well plate and blocked with 2 mM thymidine in parallel with samples used for the imaging. Then, cells were fixed with 4% formaldehyde, washed twice with PBS, stained with FxCycle PI/RNase Staining Solution (Invitrogen #F10797) and analyzed using BD Accuri C6 cytometer. Signal from more than 100,000 cells was collected and data were further processed and analyzed using FCS Express (De Novo Software).

### Single-molecule live-cell imaging

HaloTagged cell lines were plated into 24-well #1.5H glass bottom dish (Celvis #P24-1.5H-N) two days prior to imaging (40,000 cells for U2OS or 65,000 cells for HeLa cell lines). To facilitate single-molecule detection, cells were sparsely labeled with JFX650 Halo-ligand using these conditions: 1 nM for 1 min (TRF1, TIN2, TPP1, POT1) or 1 nM for 30 sec (TRF2, RAP1) in complete media at 37°C. Then, cells were washed twice with complete media and incubated for additional 15 min. After incubation, cell nuclei were stained for 2 min with Hoechst dye diluted 1:5,000 in complete media followed by two-times wash. The live-cell imaging was performed using the HILO (Highly Inclined and Laminated Optical Sheet) modality on Olympus IX83 microscope with 640 nm laser line (set to 35% power). The imaging speed was set to 100 Hz (equals to 10 ms time delay). Field of view was adjusted to 512 x 512 px (BT-Fusion camera) or 400 x 400 px with 2 x 2 px binning (Orca-Quest camera). With these settings, 1000 frames were captured for each acquisition. After each recorded movie, one image of Hoechst channel was taken using LED 385 nm light as a nuclear marker for the segmentation of single-molecule signal from nuclei. For combining the sparse and dense labeling of Halo-shelterin proteins to visualize single-molecules together with telomeric foci, the cells were first sparsely labeled with JF657 Halo-ligand using these conditions: 2 nM for 2 min (TRF1, TIN2, TPP1, POT1) or 2 nM for 1 min (TRF2, RAP1) in complete media at 37°C followed by washing cells twice with complete media and then densely labeled with 250 nM JF549 Halo-ligand for 15 min. After labeling, cells were washed three-times and incubated for additional 15 min in complete media. The live-cell imaging was performed with the same settings as above with this modification: after acquiring 1000 frames of single-molecule JF657 signal, the Z-stack using epi-illumination with 488 nm laser line was collected to mark telomeric shelterin foci. Z-stack was then maximum intensity projected and overlayed with single-molecule movies.

### Single-particle tracking (SPT) analysis

For localizing single molecules and linking them into trajectories, the multiple-target tracing algorithm(70) was used with modified parallel processing version of SLIMfast in MATLAB(54). These settings were used to track single particles: exposure time = 10 ms, NA = 1.5, pixel size = 0.1083 μm (BT-Fusion camera) or 0.1533 μm (Orca-Quest camera), emission wavelength = 667 nm (for Halo-JFX650) or 672 nm (for Halo-JF657), D_max_ = 5 μm^2^ s^−1^, number of gaps allowed = 2, localization error = 10^-6^, deflation loops = 0. Images of nuclei channel were used to generate nuclear masks based on which only nuclear trajectories were segmented and used for analysis. The MATLAB version of Spot-On algorithm(54) was employed to analyze tracks according to mean-square displacements (MSDs) to derive the diffusion coefficient of tracked single particles and respective fraction of molecules residing in either bound or freely diffusing state (2-state model was used in this analysis). The Spot-On algorithm was used with these parameters: time gap = 10 ms, dZ = 0.700 μm, gaps allowed = 2, time points = 7, jumps to consider = 4, bin width = 0.01 μm, CDF fitting routine, D_free_ 2-state boundaries = [0.5 10], D_bound_ 2-state boundaries = [0.0001 0.5]. Alternatively, trajectories were also analyzed using the ExTrack procedure that describes each trajectory as a probability function and derives underlying biophysical properties, including transition rates, based on Hidden-Markov-Modelling (HMM) (55). The ExTrack analysis was performed in Python with these settings: number of states = 2, minimal trajectory length = 3, dT = 0.01 s, frame length = 6, max distance allowed for consecutive position = 1.0, number of positions per track accepted = [3, 30], localization error type = 1, D_max_ = 5 μm^2^ s^−1^, optimization method = ‘Powell’. All experiments were carried out as three independent biological replicates with at least 30 cells for each sample. All statistical analysis were performed using GraphPad Prism by two-way ANOVA with Tukey’s posthoc test.

### Residence time imaging and analysis

HaloTagged shelterin cell lines were plated and sparsely labeled with Halo-JF657 ligand according to the protocol described in “Single-molecule live-cell imaging”. Halo-JF657 was used due to its superior photostability performance (67). Long-term single-molecule imaging was performed using HILO modality on Olympus IX83 microscope with 20% laser power (640 nm laser line) and 100 ms exposure time to blur out freely diffusing particles. Time delay between frames was set to either 500 or 1500 ms and a total of 250 frames were collected. This approach allowed to measure residence time with two distinct temporal conditions. To track only static particles, the D_max_ was set to 0.0075 μm^2^ s^−1^ and to limit the possibility of chopping long trajectories into several shorter ones, the number of gaps allowed was set to 5 and localization error to 10^-5^. Only nuclear single-particle trajectories were used for the analysis (segmented according to nuclear masks generated from Hoechst channel). Additionally, trajectories with a total length of less than 3 were not included into the analysis. Movies with 500 ms time delay were processed by averaging 3 consecutive frames to match the temporal resolution of movies with 1500 ms. The distribution of track length was visualized as a survival distribution obtained as a 1-CDF (Cumulative Density Function of track lengths) in MATLAB and fitted with two-component exponential function to extract the respective rate constants. All experiments were carried out as three independent biological replicates with at least 10 cells for each sample. All statistical analyses were performed using GraphPad Prism by two-way ANOVA with Tukey’s posthoc test.

### Conditional degradation of HaloTagged proteins for studying shelterin recruitment to telomeres

HaloTagged shelterin cell lines (40,000 U2OS and 65,000 HeLa cells) were plated into 24-well #1.5H glass bottom dish (Celvis #P24-1.5H-N). Next day morning, cells were transfected with 100 ng of plasmids coding for shelterin factors fused with mNeonGreen fluorescent protein using Lipofectamine 2000 (Invitrogen) according to the manufacturer protocol. After ∼5 h of transfection, cells were washed twice and incubated with fresh media supplemented with 500 nM Halo-PROTAC-E ligand (Aobious AOB13155) to specifically degrade the endogenous HaloTagged shelterin proteins. After ∼18 hours of incubation, cells were labeled with 250 nM Halo-JFX650 ligand for 45 min with the presence of 500 nM of Halo-PROTAC-E. Then, cells were washed twice, and nuclei were stained using Hoechst dye (1:5,000 dilution) for 4 min followed by washing cells twice and finally keeping cells in fresh complete media supplemented with 500 nM Halo-PROTAC-E ligand for imaging. Live-cell imaging was carried out using 3i spinning-disc confocal microscope with SoRa modality. Acquired Z-stacks were processed in ImageJ to obtain maximum intensity projections.

## SUPPLEMENTAL MATERIAL

**Supplementary Figure 1.**
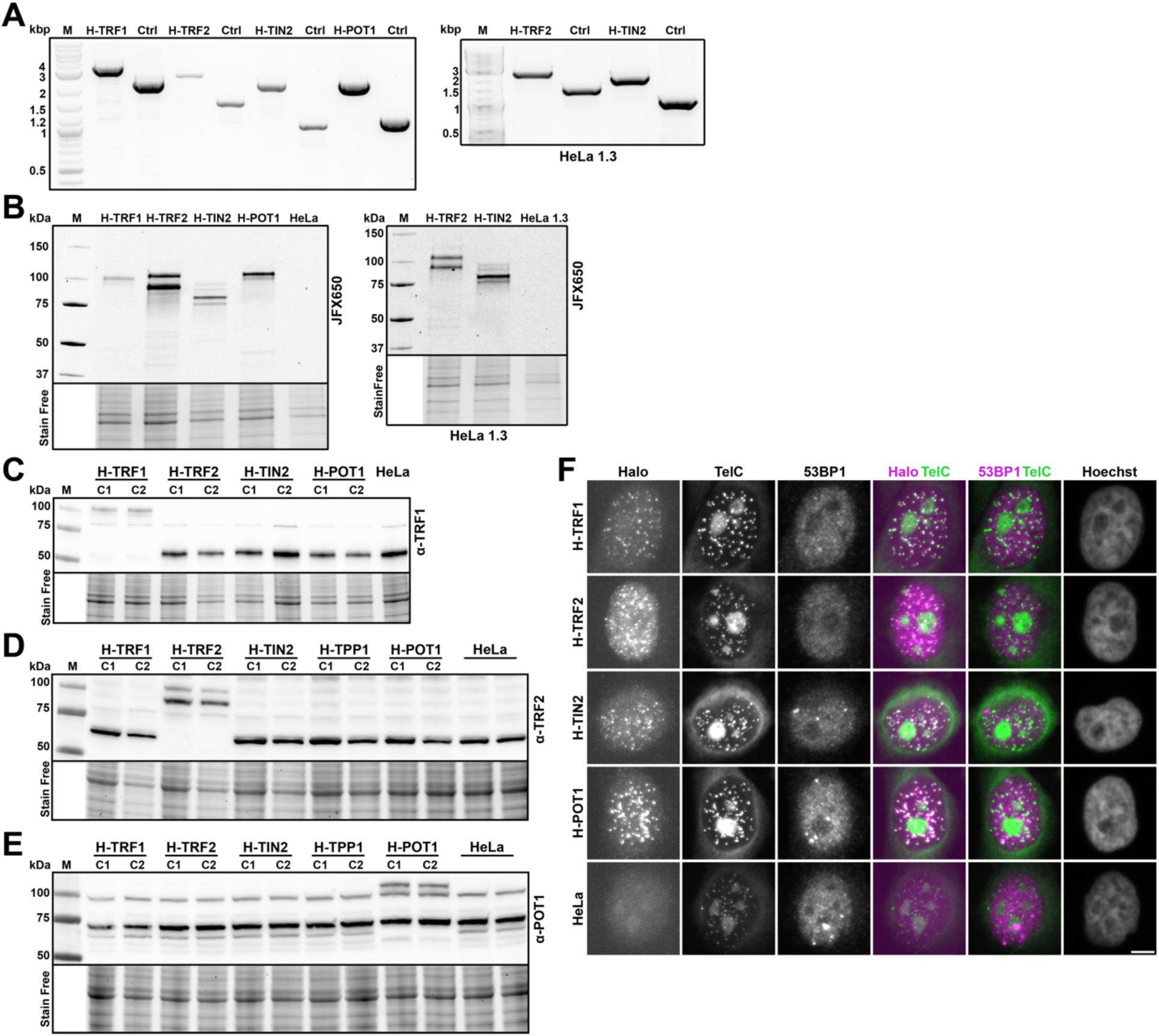
Generating a panel of HaloTagged shelterin components. **(A)** Genomic PCR result using primers outside of homology arms showing homozygous insertion of HaloTag to shelterin genes in HeLa cells (left) and HeLa 1.3 cells (right) **(B)** Representative in-gel fluorescence images demonstrating the expression of HaloTagged shelterin proteins at correct size in HeLa cells (left) and HeLa 1.3 cells (right). **(C-E)** Western blot analysis of all HeLa cell lines expressing HaloTagged shelterin proteins using antibodies against **(C)** TRF1, **(D**) TRF2, or **(E)** POT1. C1 and C2 denote clone number. **(F)** IF-FISH analysis of HeLa cell lines expressing HaloTagged shelterin proteins. The HaloTag was labeled using JFX650 HaloTag-ligand, immunofluorescence against 53BP1 was used to analyze DNA damage signaling at telomeres, and PNA FISH to mark telomeres. Scale bar = 5 µm.

**Supplementary Figure 2.**
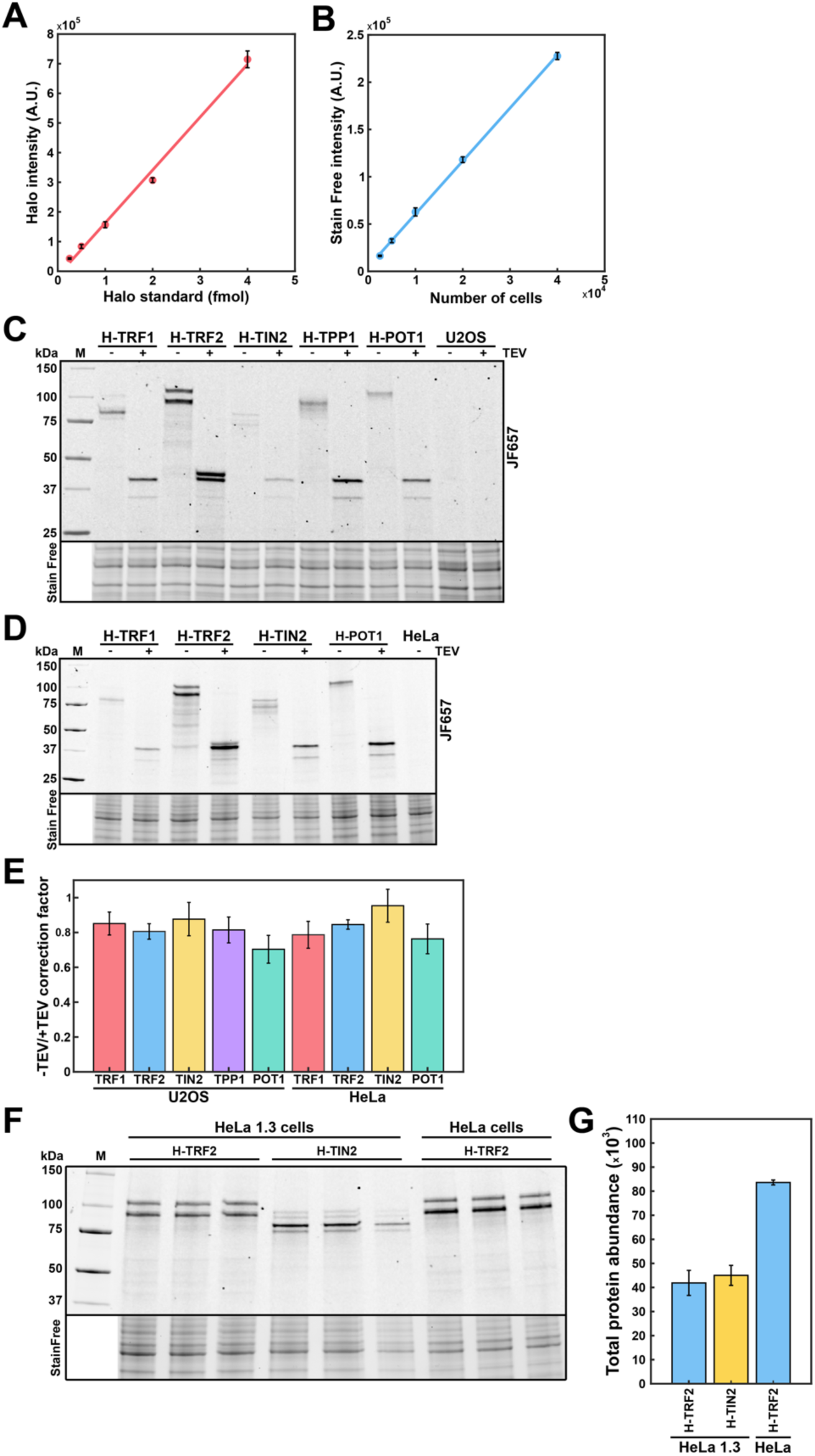
Quantification of total cellular protein abundance of HaloTagged shelterin proteins. **(A)** Standard curve for HaloTag standard fluorescence signal intensity (*N* = 3, mean ± SD). **(B)** Standard curve of stain free total protein signal (*N* = 3, mean ± SD). **(C)** In-gel fluorescence of cell lysates generated from U2OS cells expressing HaloTagged shelterin proteins treated with or without TEV protease. **(D)** In-gel fluorescence of cell lysates generated from HeLa cells expressing HaloTagged shelterin proteins treated with or without TEV protease. **(E)** Quantification of the ratio of the HaloTag fluorescence intensity in untreated samples relative to TEV treated lysates from cell lines expressing HaloTagged shelterin proteins (*N* = 3, mean ± SD). **(F)** Representative in-gel fluorescence image used for the quantification of Halo-TRF2 and Halo-TIN2 total cellular abundance in HeLa 1.3 cells compared to the signal of Halo-TRF2 from HeLa cells. Image also demonstrates the expression of HaloTagged shelterin proteins at correct size in HeLa 1.3 cells. **(G)** Quantification of the total cellular abundance of Halo-TRF2 and Halo-TIN2 in HeLa 1.3 cells. Data was generated by comparing the fluorescence intensity of Halo-TRF2 and Halo-TIN2 in HeLa 1.3 cells to Halo-TRF2 in HeLa cells, normalized using the total protein stain signal from (F).

**Supplementary Figure 3.**
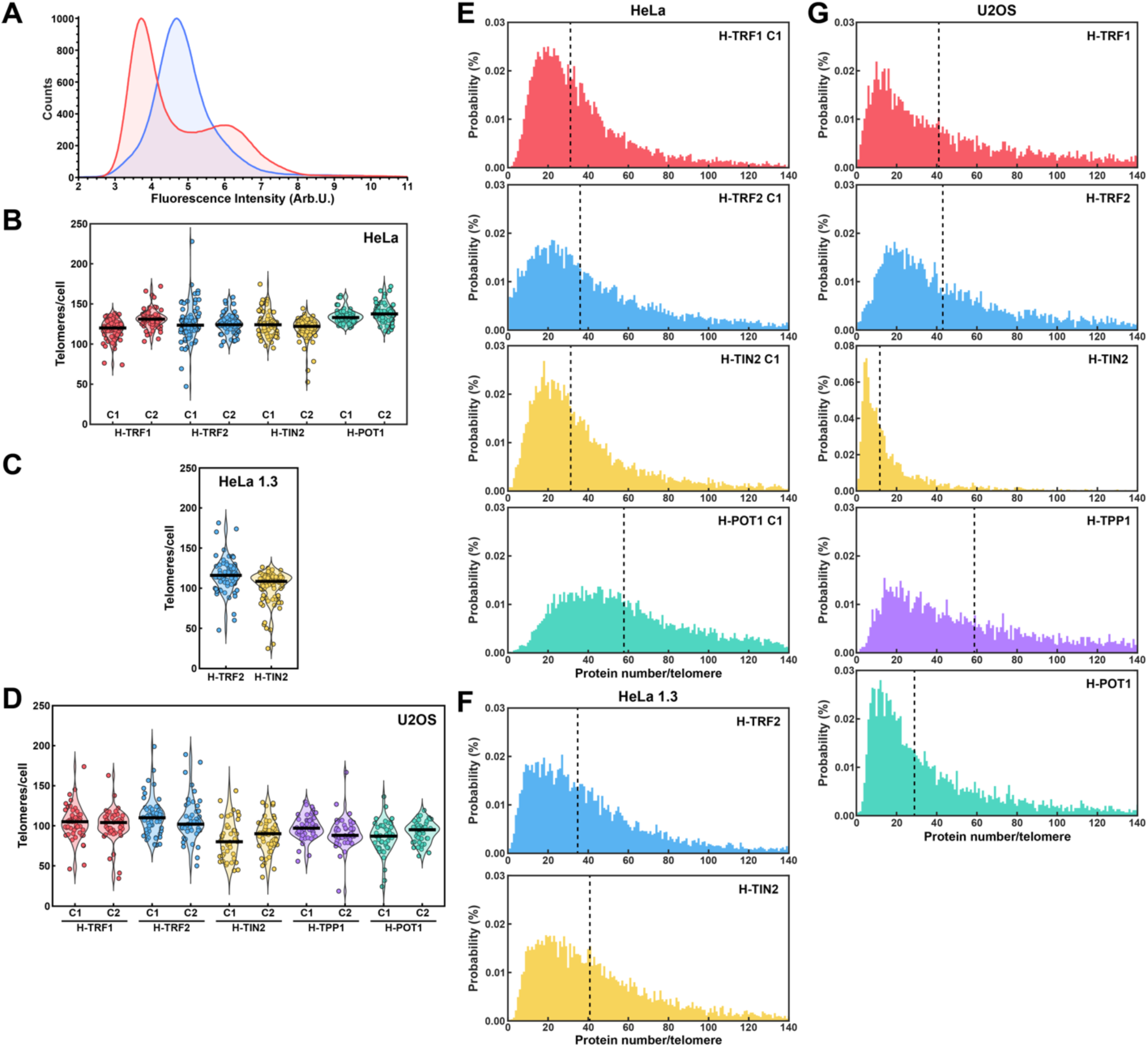
Quantification of telomeric protein number of HaloTagged shelterin proteins. **(A)** DNA content analysis using flow cytometry with propidium iodide staining of asynchronous HeLa cells and HeLa cells synchronized using a double thymidine block. **(B-D)** Quantification of the number of nuclear foci formed by the HaloTagged shelterin proteins in **(B)** HeLa, **(C)** HeLa 1.3, and **(D)** U2OS cells. Analyzed images were maximum intensity projections of 51 Z-sections from a single biological replicate used for the quantification of telomeric protein number of HaloTagged shelterin proteins. **(E-G)** Distributions of telomeric abundance of HaloTagged shelterin proteins from a single biological replicate generated from **(E)** HeLa, **(F)** HeLa 1.3, and **(G)** U2OS cells (At least 60 cells analyzed per cell line, dashed line indicates median telomeric abundance).

**Supplementary Figure 4.**
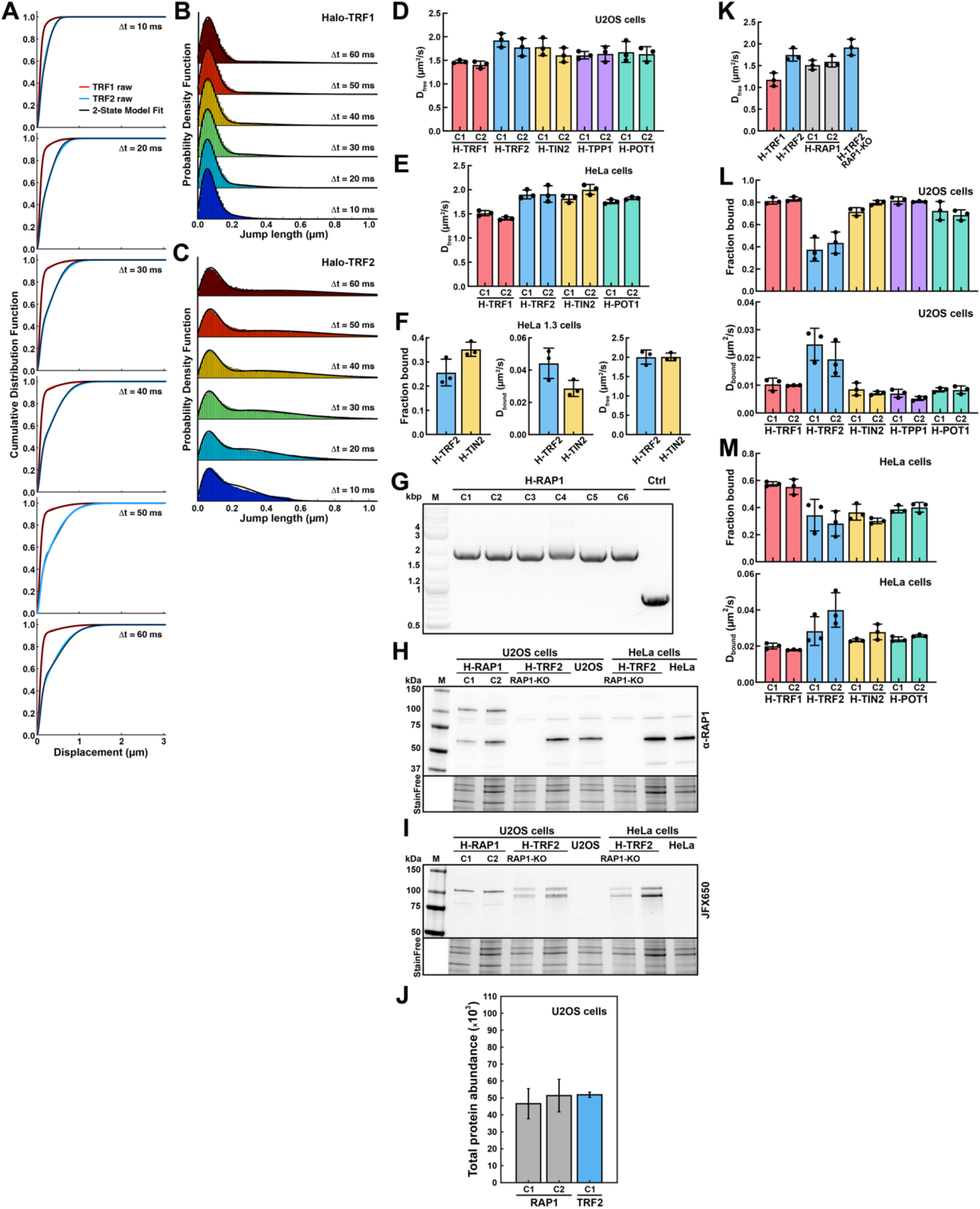
Single-molecule analysis of shelterin dynamics. **(A)** Cumulative distribution function of the displacements of single-particle trajectories generated from Halo-TRF1 (red) and Halo-TRF2 (blue) expressing U2OS cells at different time delays. Colored lines represent the raw data, and black lines indicate the Spot-On fit assuming a two-state model with a static and freely diffusing population. **(B-C)** Probability density function of single-particle jump sizes generated from **(B)** Halo-TRF1 and **(C)** Halo-TRF2 expressing U2OS cells at different time delays. Colored bars indicate raw data, black lines indicate the model fit generated from the cumulative distribution function analysis (Figure S4A). **(D-E)** Quantification of the diffusion coefficient of freely diffusing HaloTagged shelterin components in **(D)** U2OS cells and **(E)** HeLa cells derived from a two-state fit of the cumulative distribution function of the single-particle displacements using the Spot-On software (*N* = 3 biological replicates, at least 30 cells per biological replicate, mean ± SD). **(F)** Quantification of the fraction of static molecules (left) and the diffusion coefficients of static (middle) and freely diffusion (right) Halo-TRF2 and Halo-TIN2 molecules in HeLa 1.3 cells using the Spot-On tool (*N* = 3 biological replicates, at least 30 cells per biological replicate, mean ± SD). **(G)** Genomic PCR using primers outside of homology arms of Halo-RAP1 clones (U2OS cells). **(H)** Western blot using an anti-RAP1 antibody demonstrating RAP1 knock-out in U2OS and HeLa cells together with the expression of Halo-RAP1 in U2OS cells. **(I)** In-gel fluorescence demonstrating the expression of Halo-RAP1 in U2OS cells and Halo-TRF2 with RAP1 knock-out in U2OS and HeLa cells. Same gel that was used for the WB in (H). **(J)** Quantification of total protein abundance of Halo-RAP1 in U2OS cells. The intensity of the Halo-RAP1 band was compared to Halo-TRF2 and normalized to total protein stain. **(K)** Quantification of the diffusion coefficient of freely diffusing Halo-TRF1, Halo-TRF2, Halo-RAP1, and Halo-TRF2 in RAP1 knock-out U2OS cells derived from a two-state fit of the cumulative distribution function of the single-particle displacements using the Spot-On software (*N* = 3 biological replicates, at least 30 cells per biological replicate, mean ± SD). **(L-M)** Quantification of the fraction of static molecules and the diffusion coefficient of static HaloTagged shelterin protein expressed in **(L)** U2OS and **(M)** HeLa cells using the ExTrack tool to analyze the same data shown in Figure 4 (*N* = 3 biological replicates, at least 30 cells per biological replicate, mean ± SD).

**Supplementary Figure 5.**
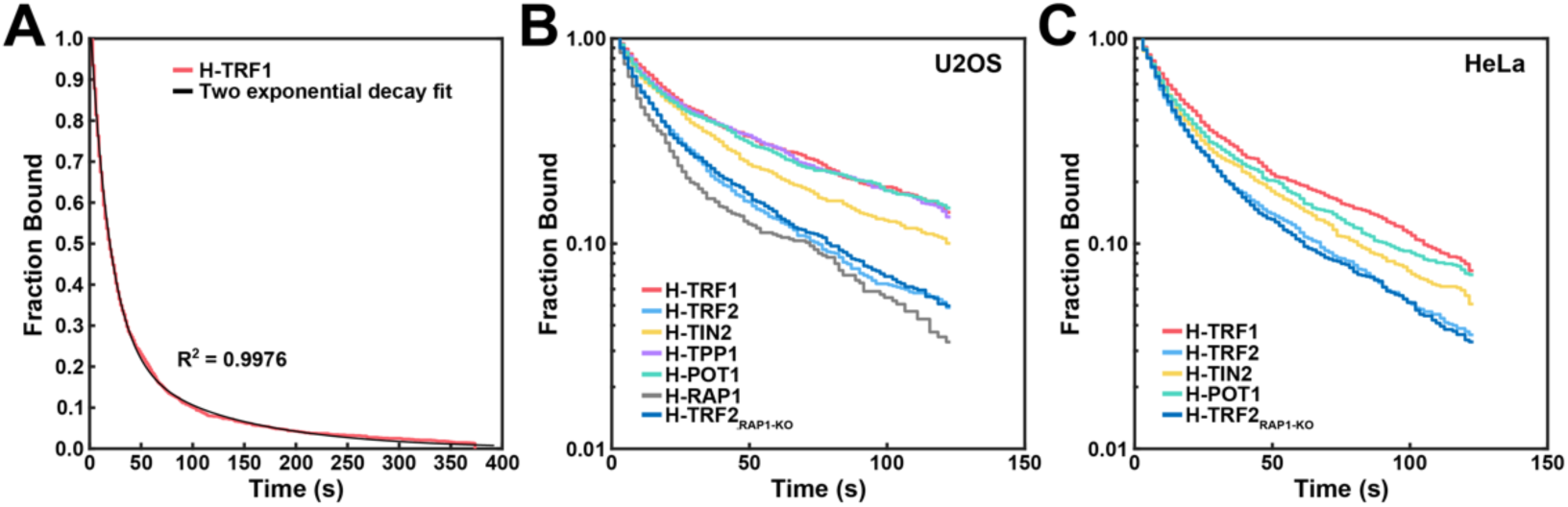
Residence time analysis of HaloTagged shelterin proteins at telomeres. **(A)** Survival distribution (red) of static Halo-TRF1 molecules generated by single particle tracking of movies acquired at 1.5 second time intervals. The black line indicates the fit of the data using two exponential decay function. **(B-C)** Representative survival distributions of single biological replicate of static shelterin molecules in **(B)** U2OS cells and **(C)** HeLa cells generated by single-particle tracking of immobile single-molecule signals of movies acquired at 0.5 second time intervals with a subsequent averaging of three consecutive frames to match the 1.5 s time interval.

**Supplementary Figure 6.**
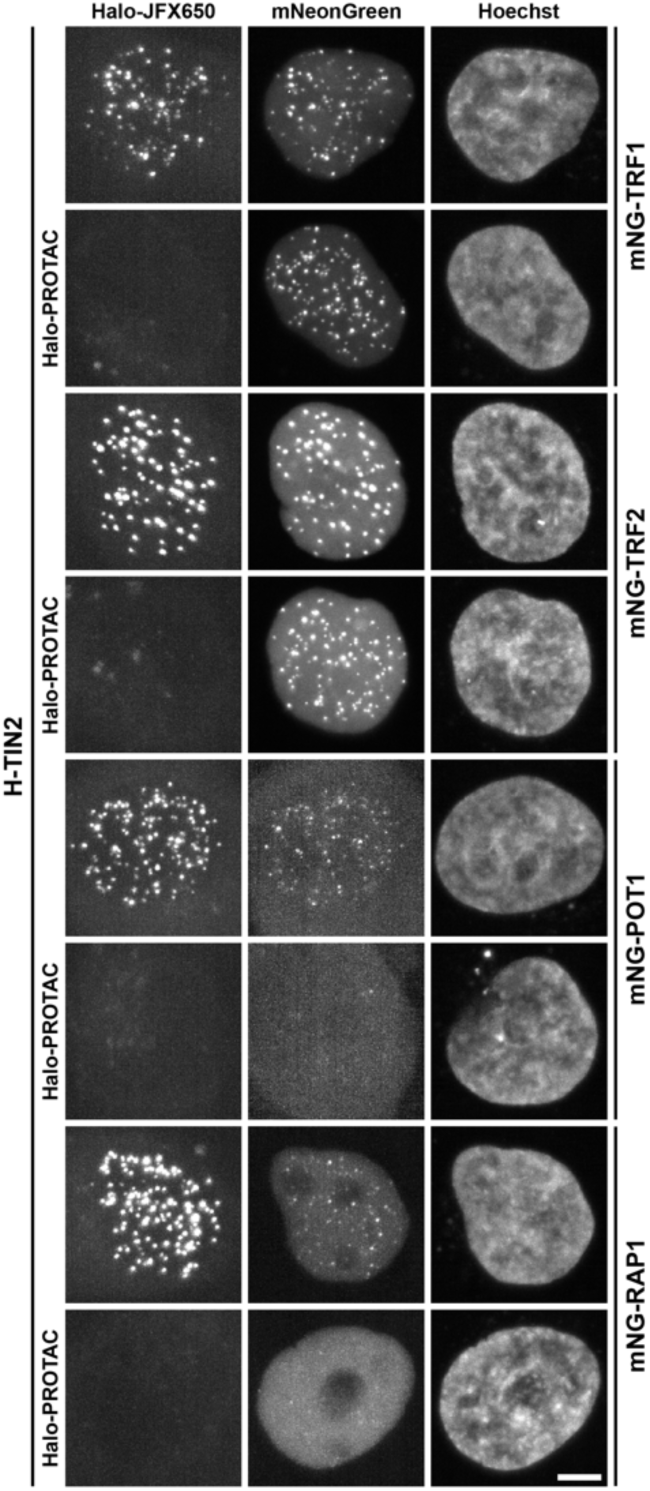
Recruitment of shelterin to telomeres. Representative images of Halo-TIN2 HeLa cells transiently expressing mNeonGreen fusion proteins of TRF1, TRF2, POT1 and RAP1, untreated or treated with Halo-PROTAC for 18 hours. Cells were labeled with JFX650 HaloTag-ligand immediately before live-cell imaging. Scale bar = 5 µm.

## SUPPLEMENTAL MOVIE LEGENDS

**Movie S1.** Movie of Halo-TRF1 expressed in U2OS cells for fluorescence photobleaching analysis to determine the fluorescence intensity of a single HaloTag labeled with JFX650 HaloTag-ligand. Acquired using a Hamamatsu ORCA-Quest camera without binning at 25 frames per second.

**Movie S2.** Single-particle tracking of Halo-TRF2 expressed in HeLa cells labeled with JFX650 HaloTag ligand. Acquired using a Hamamatsu ORCA-Fusion BT camera at 100 frames per second.

**Movie S3.** Live-cell single-molecule imaging movie of Halo-TRF1 expressed in U2OS cells labeled with JFX650 HaloTag ligand. Acquired using a Hamamatsu ORCA-Fusion BT camera at 100 frames per second.

**Movie S4.** Live-cell single-molecule imaging movie of Halo-TRF2 expressed in U2OS cells labeled with JFX650 HaloTag ligand. Acquired using a Hamamatsu ORCA-Fusion BT camera at 100 frames per second.

**Movie S5.** Live-cell single-molecule imaging movie of Halo-TIN2 expressed in U2OS cells labeled with JFX650 HaloTag ligand. Acquired using a Hamamatsu ORCA-Fusion BT camera at 100 frames per second.

**Movie S6.** Live-cell single-molecule imaging movie of Halo-TPP1 expressed in U2OS cells labeled with JFX650 HaloTag ligand. Acquired using a Hamamatsu ORCA-Fusion BT camera at 100 frames per second.

**Movie S7.** Live-cell single-molecule imaging movie of Halo-POT1 expressed in U2OS cells labeled with JFX650 HaloTag ligand. Acquired using a Hamamatsu ORCA-Fusion BT camera at 100 frames per second.

**Movie S8.** Live-cell single-molecule imaging movie of Halo-TRF1 expressed in HeLa cells labeled with JFX650 HaloTag ligand. Acquired using a Hamamatsu ORCA-Fusion BT camera at 100 frames per second.

**Movie S9.** Live-cell single-molecule imaging movie of Halo-TRF2 expressed in HeLa cells labeled with JFX650 HaloTag ligand. Acquired using a Hamamatsu ORCA-Fusion BT camera at 100 frames per second.

**Movie S10.** Live-cell single-molecule imaging movie of Halo-TIN2 expressed in HeLa cells labeled with JFX650 HaloTag ligand. Acquired using a Hamamatsu ORCA-Fusion BT camera at 100 frames per second.

**Movie S11.** Live-cell single-molecule imaging movie of Halo-POT1 expressed in HeLa cells labeled with JFX650 HaloTag ligand. Acquired using a Hamamatsu ORCA-Fusion BT camera at 100 frames per second.

**Movie S12.** Live-cell single-molecule imaging movie of Halo-TRF2 expressed in HeLa 1.3 cells labeled with JFX650 HaloTag ligand. Acquired using a Hamamatsu ORCA-Quest camera using 2 x 2 px binning at 100 frames per second.

**Movie S13.** Live-cell single-molecule imaging movie of Halo-TIN2 expressed in HeLa 1.3 cells labeled with JFX650 HaloTag ligand. Acquired using a Hamamatsu ORCA-Quest camera using 2 x 2 px binning at 100 frames per second.

**Movie S14.** Dual-color live-cell single-molecule imaging of Halo-TRF1 expressed in HeLa cells sparsely labeled with JFX650 HaloTag-ligand for single-molecule detection, followed by quantitative labeling with JF549 HaloTag-ligand to detect telomeric foci. Telomeric foci were imaged as a Z-stack after single-molecule acquisition. A maximum intensity projection of the telomeric foci was overlayed with the single-molecule movie. Acquired using a Hamamatsu ORCA-Quest camera using 2 x 2 px binning at 100 frames per second.

**Movie S15.** Dual-color live-cell single-molecule imaging of Halo-TRF2 expressed in HeLa cells sparsely labeled with JFX650 HaloTag-ligand for single-molecule detection, followed by quantitative labeling with JF549 HaloTag-ligand to detect telomeric foci. Telomeric foci were imaged as a Z-stack after single-molecule acquisition. A maximum intensity projection of the telomeric foci was overlayed with the single-molecule movie. Acquired using a Hamamatsu ORCA QUEST camera using 2x2 binning at 100 frames per second.

**Movie S16.** Dual-color live-cell single-molecule imaging of Halo-TIN2 expressed in HeLa cells sparsely labeled with JFX650 HaloTag-ligand for single-molecule detection, followed by quantitative labeling with JF549 HaloTag-ligand to detect telomeric foci. Telomeric foci were imaged as a Z-stack after single-molecule acquisition. A maximum intensity projection of the telomeric foci was overlayed with the single-molecule movie. Acquired using a Hamamatsu ORCA-Quest camera using 2 x 2 px binning at 100 frames per second.

**Movie S17.** Dual-color live-cell single-molecule imaging of Halo-POT1 expressed in HeLa cells sparsely labeled with JFX650 HaloTag-ligand for single-molecule detection, followed by quantitative labeling with JF549 HaloTag-ligand to detect telomeric foci. Telomeric foci were imaged as a Z-stack after single-molecule acquisition. A maximum intensity projection of the telomeric foci was overlayed with the single-molecule movie. Acquired using a Hamamatsu ORCA-Quest camera using 2 x 2 px binning at 100 frames per second.

**Movie S18.** Live-cell single-molecule imaging movie of Halo-RAP1 expressed in U2OS cells labeled with JFX650 HaloTag ligand. Acquired using a Hamamatsu ORCA-Quest camera using 2 x 2 px binning at 100 frames per second.

**Movie S19.** Live-cell single-molecule imaging movie of Halo-TRF2 expressed in U2OS cells with a RAP1 knock-out labeled with JFX650 HaloTag ligand. Acquired using a Hamamatsu ORCA-Quest camera using 2 x 2 px binning at 100 frames per second.

**Movie S20.** Live-cell single-molecule imaging movie of Halo-TRF1 expressed in HeLa cells labeled with JFX650 HaloTag ligand. Acquired using a Hamamatsu ORCA-Quest camera using 2 x 2 px binning at 1 frame every 1.5 seconds.

## Notes

### Competing Interest Statement

The authors have declared no competing interest.

